# Efficacy of *Aedes aegypti* control by indoor Ultra Low Volume (ULV) insecticide spraying in Iquitos, Peru

**DOI:** 10.1101/231134

**Authors:** Christian Gunning, Kenichi Okamoto, Helvio Astete, Gissella M. Vasquez, Erik Erhardt, Clara Del Aguila, Raul Pinedo, Roldan Cardenas, Carlos Pacheco, Enrique Chalco, Hugo Rodriguez-Ferruci, Thomas W. Scott, Alun L. Lloyd, Fred Gould, Amy C. Morrison

## Abstract

**Background:** *Aedes aegypti* is a primary vector of dengue, chikungunya, Zika, and urban yellow fever viruses. Indoor, ultra low volume (ULV) space spraying with pyrethroid insecticides is the main approach used for *Ae. aegypti* emergency control in many countries. Given the widespread use of this method, the lack of large-scale experiments or detailed evaluations of municipal spray programs is problematic.

**Methodology/Principal Findings:** Two experimental evaluations of non-residual, indoor ULV pyrethroid spraying were conducted in Iquitos, Peru. In each, a central sprayed sector was surrounded by an unsprayed buffer sector. In 2013, spray and buffer sectors included 398 and 765 houses, respectively. Spraying reduced the mean number of adults captured per house by ~83 percent relative to the pre-spray baseline survey. In the 2014 experiment, sprayed and buffer sectors included 1,117 and 1,049 houses, respectively. Here, the sprayed sector’s number of adults per house was reduced ~64 percent relative to baseline. Parity surveys in the sprayed sector during the 2014 spray period indicated an increase in the proportion of very young females. We also evaluated impacts of a 2014 citywide spray program by the local Ministry of Health, which reduced adult populations by ~60 percent. In all cases, adult densities returned to near-baseline levels within one month.

**Conclusions/Significance:** Our results demonstrate that densities of adult *Ae. aegypti* can be reduced by experimental and municipal spraying programs. The finding that adult densities return to approximately pre-spray densities in less than a month is similar to results from previous, smaller scale experiments. Our results demonstrate that ULV spraying is best viewed as having a short-term entomological effect. The epidemiological impact of ULV spraying will need evaluation in future trials that measure capacity of insecticide spraying to reduce disease transmission.

## INTRODUCTION

*Aedes aegypti* is a primary vector for dengue (DENV), chikungunya (CHIKV), Zika (ZIKV) and urban yellow fever viruses (YFV). Dengue has become the most important human arthropod-borne viral infection worldwide (Brady et al. 2012, Bhatt et al. 2013). Each of these pathogens can be associated with explosive epidemics, where high disease incidence and public fear combine to overwhelm health systems (Wilder-Smith et al. 2016). Such epidemics put intense pressure on public health departments to react with emergency vector control measures (Esu et al. 2010, Simmons et al. 2012).

*Ae. aegypti* adults are primarily diurnal and females take frequent blood meals, predominantly from humans (Scott et al 1997, 2000, Scott & Takken 2012). These behaviors can in part explain why *Ae. aegypti* has been associated with epidemic virus transmission even when its population densities are low (Kuno 1995). Because adults typically reside inside houses (Scott & Takken 2012) where food, mates, and oviposition substrates are readily available, indoor adulticide space spraying has been more effective than outdoor spraying for suppressing *Ae. aegypti* populations (Morrison et al. 2008, Reiter et al. 2014, Esu et al 2010).

When indoor space sprays are applied appropriately, in carefully controlled small-scale expermants, adult *Ae. aegypti* populations often decreased by >80%. Population densities typically recovered quickly, however, (Perich et al. 2000, 2001, 2003; Koenraadt et al. 2007; Bowman et al. 2016) due to emergence of nulliparous mosquitoes from larval aquatic habitats inside sprayed areas (Reiter 2014), through migration from locations outside of sprayed areas (Koenraadt et al. 2007), or from females in sprayed houses that survived. In a systematic literature review, Esu et al. (2010) found only six studies from 1970’s to 2010 that tested ultra-low volume (ULV) indoor space spraying under natural field conditions that met minimum standards for evaluating mosquito population suppression. None of the studies evaluated the impact of these methods on human infection or disease (Esu et al. 2010). Results ranged from immediate reduction in biting by 99% and adult population reduction lasting six months (Pant et al. 1974), to a more common, modest control lasting 1-5 weeks (Perich et al. 2001; Koenraadt et al. 2007, Castro et al. 2007). Most studies were small scale, with each treatment typically including one replicate of less than 50 houses. A more recent review of vector control effectiveness for dengue (Bowman et al. 2016) concluded that “although space spraying is the standard public health response to a dengue outbreak worldwide, and is recommended by WHO (2011) for this purpose, there is scant evidence available from studies to evaluate this method sufficiently.” In fact, Bowman et al. [26] (2016) could find no well-designed trial that assessed the impact of non-residual space spraying on human dengue infection or disease.

*Ae. aegypti* populations in the Amazonian city of Iquitos, Peru have been studied extensively since 1998. The spatial distribution of the species is highly clustered and does not have a consistent spatial or temporal structure (Getis et al. 2003, LeCon et al. 2014). Adult and immature population indices are highly variable and subject to sampling error (Morrison et al. 2004a). Evaluation of control measures for this species, therefore, requires large sample sizes and exhaustive sampling.

In addition to studying the mosquito itself, the Iquitos research program monitored dengue transmission through passive clinic-based febrile surveillance in health care facilities throughout the city (Forshey et al. 2010) and a series of prospective cohort studies in targeted city neighborhoods (Morrison et al 2010, Rocha et al 2009, Stoddard et al 2013). The combination of longitudinal entomological and epidemiological studies created a database that could be used to examine, in real time, the impact of Ministry of Health (MoH) vector interventions on *Ae. aegypti* populations and human disease. During their interventions, the MoH sprayed non-residual insecticide inside homes three times over an approximately 3-week period (Stoddard et al. 2014). Over a 10-year period, this kind of citywide municipal vector control program was associated with significant decreases in *Ae. aegypti* adult populations (Morrison et al. 2003, 2005) and when interventions were applied during the first half of the dengue transmission season, fewer dengue cases were detected and the transmission season was shorter (Stoddard et al. 2014). While the qualitative results from that analysis of dengue are consistent with an expectation of a positive public health impact of intra-domicile ULV insecticide application on dengue incidence, more statistically robust epidemiological studies are needed (Reiner et al. 2016).

Prevention of *Aedes*-transmitted viral disease will require integrated approaches; i.e., combinations of existing and/or novel vector control strategies as well as vaccination. Mathematical models provide a way to compare diverse strategies and identify the most promising approaches. For example, data on *Ae. aegypti* populations in Iquitos were used to develop a biologically detailed, spatially explicit, stochastic model that tracked *Ae. aegypti* dynamics and genetics in an 18-ha area of the city (Legros et al. 2011, Magori et al. 2009). Preliminary validation of the model using Iquitos data was carried out (Legros et al. 2011), but evaluation of its capacity to accurately predict the entomological outcome of a vector control perturbation had not been tested. The experiments described here were primarily designed to generate data that could be used to test the ability of the entomological model to predict impacts of suppression measures.

In this study, we carried out a large-scale evaluation of the entomological impact of a widely used emergency vector intervention of Aedes-transmitted viruses in a well-characterized study site. Our specific goal was to evaluate the impact of 6 cycles of indoor ULV pyrethroid spray applications (hereafter referred to as “spray applications”) on reductions of *Ae. aegypti* populations. Our experiments spanned periods of relatively low and high *Ae. aegypti* density in Iquitos, and compared the ULV application in experimental and public health settings. Our results constitute an important data set for development and validation of *Ae. aegypti* population dynamics models, and provide a detailed account of indoor space spray effects on *Ae. aegypti* populations.

## METHODS AND MATERIALS

### Study Area

Our studies were conducted in two neighborhoods in the Maynas district of Iquitos (Fig. 1, Maps). Iquitos has a human population of ~380,000 (73.2’W longitude, 3.7°S latitude, 120 m above sea level). Located in the Amazon Basin of northeastern Peru, Iquitos is the largest urban center in the Department of Loreto, and has an average daily temperature of 25°C and an average annual precipitation of 2.7 meters. Dynamics of *Ae. aegypti* populations in Iquitos are described in detail in earlier publications (Getis et al. 2003; Morrison et al. 2004a,b, 2006, 2010; LeCon et al. 2014; Stoddard et al. 2013; Schneider et al. 2004, Hayes et al 1996; Watts et al. 199; Paz-Soldan 2011)

**Figure 1.**
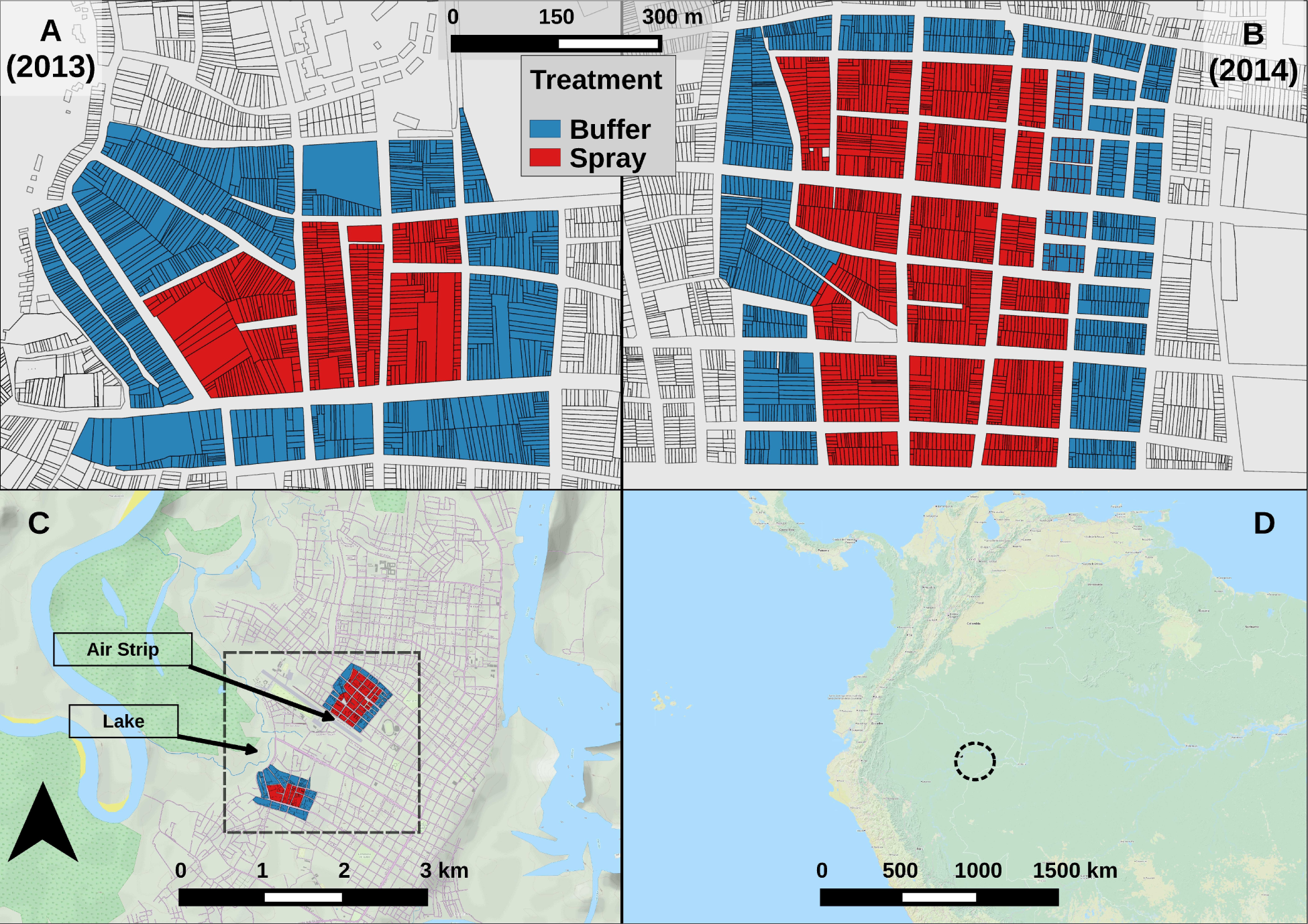
Map of experiment areas. **A, B**: Detail of experimental areas, showing individual houses. Color shows sector. **C**: City of Iquitos. Black box highlights experimental areas. **D**: Regional map. Black circle highlights Iquitos. See also Fig. S6.

Both experimental study neighborhoods were characterized by city blocks of row houses (dwellings that share walls). Most houses occupied lots that were narrow (3-10 m wide), but relatively deep (20-60 m long). The majority of houses served as family residences, often containing extended or multiple families. Some houses were used for small businesses or offices, and others were unoccupied. There were a small number of vacant lots containing no structures (<1%). Many study houses were mixed-purpose, sharing living areas with a small store (“bodega”), office, shop (e.g. carpentry or vehicle repair), or restaurant.

Vector control activities were ongoing in Iquitos. The MoH carried out regular entomological surveillance and larviciding activities with temephos (®Abate) at ~3 month intervals. Since 2002, with few exceptions, MoH carried out 1-3 emergency indoor pyrethroid spray campaigns per year in response to dengue outbreaks, with variable success (Stoddard et al. 2014). Our study was completed in 2014, and resistance bioassay profiles prior to January 2015 indicated *Ae. aegypti* populations in the city were susceptible to pyrethroids (Palomino-Salcedo 2014).

Figure 2 (Flow Chart) summarizes the design of our two separate experiments. The first and smaller of the two experiments (S-2013) ran for 16 calendar weeks and included an experimental buffer sector that was not sprayed, surrounding a central experimental sector that was sprayed. The buffer sector contained 765 houses and the spray sector had 398 houses (Fig. 1A, Table 1). The S-2013 study area was located on the western border of the city, proximal to Lake Moronacocha (Fig. 1C).

**Figure 2.**
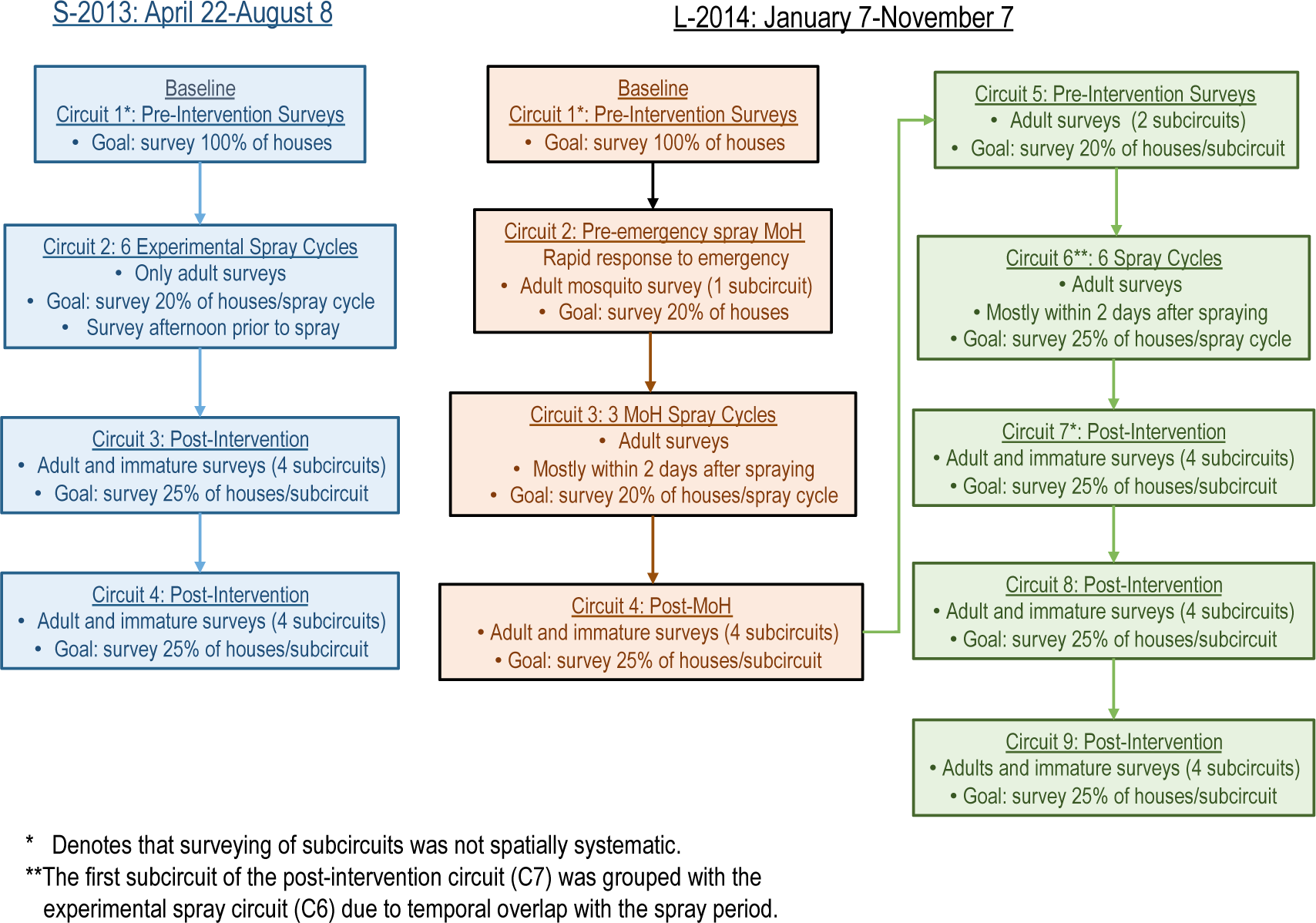
Experiment timeline. Each box shows one circuit. With one exception (L-2014 C2), each house was visited (and possibly surveyed) at least once per circuit. Except where noted, each circuit consisted of one or more spatially systematic subcircuits. Each subcircuit lasted approximately one calendar week. See Fig. S7 for survey maps.

**Table 1. [tab_count].** Observation counts, including houses, surveys, adults, containers, and adult female dissections (parity). Note that houses were surveyed repeatedly. Only *Ae. aegypti* mosquitoes are included here. Positive containers have visible eggs, larvae, or pupae. Nearly all sampled adult females (column 5) were dissected to determine parity status (columns 9 & 10). See also Table S1.

| Experiment | Sector | Houses | Surveys | Sampled | Adults: | Sampled | Containers: | Dissected | Females: |
| --- | --- | --- | --- | --- | --- | --- | --- | --- | --- |
|  |  |  |  | Female | Total | Positive | Total | Nulliparous | Total |
| S-2013 | Buffer | 765 | 2448 | 439 | 904 | 236 | 5311 | 49 | 406 |
| S-2013 | Spray | 398 | 1395 | 153 | 354 | 109 | 2170 | 23 | 142 |
| L-2014 | Buffer | 1051 | 5810 | 1585 | 3165 | 251 | 6811 | 81 | 1444 |
| L-2014 | Spray | 1110 | 6314 | 2092 | 4244 | 278 | 7454 | 191 | 2004 |

The larger second experiment (L-2014) ran for 44 calendar weeks, and included 1,051 houses in the surrounding buffer sector and 1,110 houses in the central spray sector (Fig. 1B, Table 1). L-2014 was carried out in a neighborhood several kilometers to the north of S-2013, centrally located in Iquitos, and bordered on the south by an abandoned airstrip (Fig. 1C). The L-2014 study area was selected because the *Ae. aegypti-free* airstrip provided a physical barrier to *Ae. aegypti* dispersal on one of its four sides. This experimental structure of L-2014 was selected to test our mathematical model’s ability to capture any spatial features of the recovering mosquito population.

### Entomological Surveys

To monitor population densities and age structure of *Ae. aegypti* populations, we carried out standardized adult mosquito collections using Prokopackm aspirators (Vazquez-Prokopec et al. 2009) (henceforth adult surveys) and standardized larval/pupal demographic surveys [47-49] (Focks et al. 1993, 1997, 2000) (henceforth immature surveys), except when noted. Survey protocols are described in detail in previous publications [6] (Getis et al. 2003; Morrison et al. 2004a; LaCon et al. 2014; Schneider et al. 2004, Vazquez-Prokopec et al. 2009).

Collected adults were immediately transported to a field laboratory in Iquitos for processing as described in Morrison et al. (Morrison et al. 2004b). Adult mosquitoes were sedated by cold (4°C), identified, counted, and females separated. In most cases, we scored female *Ae. aegypti* as unfed, blood fed (full, half full, or trace amounts), or gravid. Females were also scored for parity (Scott et al. 2000).

### Pyrethroid spray applications

Experimental insecticide spraying was done by MoH employees, between 17:00-20:00 to avoid high temperatures and varying winds. Each spray team was comprised of 3 individuals: 2 MoH sprayers and 1 monitor from the research team. Each week, on the initial day of a spray cycle (usually Mondays), spraying was attempted in all houses in the spray sector. To improve spray coverage within each cycle, on subsequent days spray teams revisited houses that were not sprayed on the initial day of the spray cycle (a minimum of 2 and up to 10 visits, as needed) to conduct spraying. Pyrethroid insecticides were applied using Solo or Stihl backpack sprayers with settings adjusted for ULV application, or Colt hand-held ULV sprayers. Residents were instructed not to return to their houses for a minimum of 1 hour. See Text S1 for more details.

### Quality Control for Spray Applications

As a quality control measure, for each spray cycle, 3 to 7 houses were selected to monitor efficacy of the insecticide spray. Operators did not know which houses would be selected for monitoring. For each monitored house, just after the spray operator had finished the application, a single screen cage containing adult mosquitoes was placed in each of the following locations: bedroom, living room, kitchen, and yard, based on standard WHO protocols (WHO 2005, Reiter & Nathan 2003). Each cage contained 25 adult *Ae. aegypti* of age 24-36 hours from a pathogen-free laboratory colony (Reiter et al. 2003, WHO guidelines). A separate laboratory colony was initiated for each experiment from mosquitoes collected from houses in Iquitos and held for 1-2 generations prior to use. One hour after spraying, all cages were retrieved and evaluated for knockdown (no movement), stored in a styrofoam cooler with moist paper towels for 24 hours, and then examined for mortality. When mortality was < 80%, equipment was recalibrated to ensure proper spray function on subsequent days.

*Droplet size.* Teflon treated slides were placed in 2 randomly selected houses during each spray cycle and retrieved 1 hour post-spray. Droplet size was measured using a micrometer in Motic Images Plus 2.2. Droplets were counted and measured in a 1 cm^2^ square.

### Experimental Design

Experimental study sectors are depicted in Figures 1A & B. The temporal sampling units are referred to as “circuits” because they were time periods when we completed full survey routes through all of the blocks of houses in the spray and buffer sectors (see Fig. 2 for a flow chart of experimental design, and Fig. S7 for survey maps). During each circuit, we attempted to visit and survey 100% of the houses in the entire study area at least once (with one exception, L-2014 C2). The percentage of total houses successfully surveyed and/or sprayed in each circuit ranged from 67-90%, due to closed or unoccupied houses, or residents who chose not to participate in the study (see Fig. 3B, Table S1).

**Figure 3. [ts_survey].**
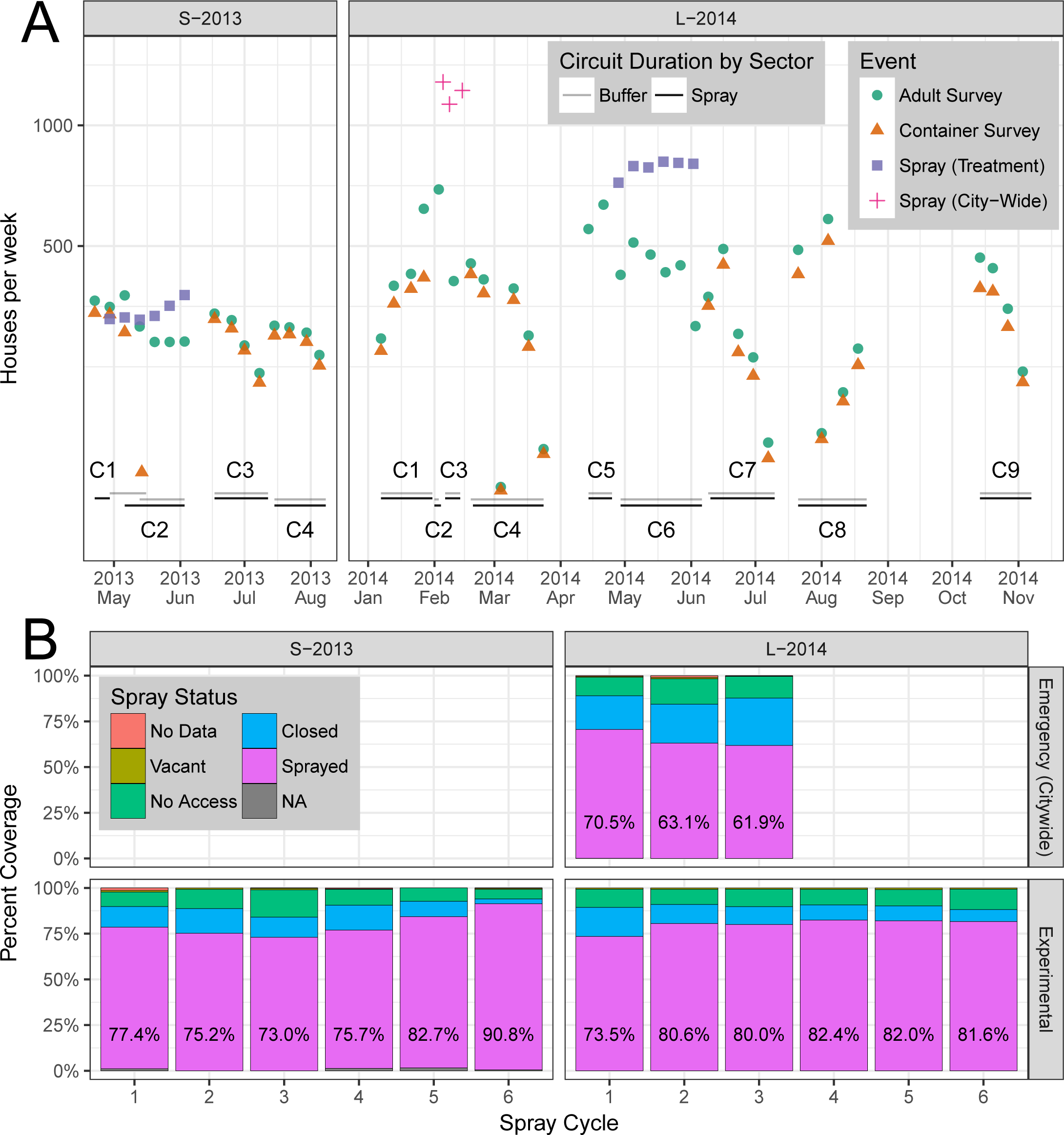
Sampling and Spraying. **A**: Number of houses per week sprayed and/or surveyed. Circuits are labeled (e.g., C1), with date ranges shown by horizontal bars. Containers were not surveyed during spray periods. The first two emergency (citywide) spray events (red +) occurred within the same calendar week, but are plotted separately here. **B**: Spray coverage by spray cycle. Percent houses sprayed is shown in text. Top row: emergency (citywide) spraying. Bottom row: experimental spraying. Note that emergency citywide spraying (3 cycles) occurred only during L-2014.

Each circuit was divided into subcircuits that lasted approximately one week, but never more than 10 days. In general, subcircuit surveying was conducted systematically by block, where surveyors attempting to visit every 4^th^ house (25% of the circuit) each week (see Text S2 for exceptions).

Both experiments consisted of 6 weekly cycles of ULV indoor spray applications (see above). Immature and adult surveys were carried out before (pre-intervention) and after (post-intervention) the spraying periods. During the experimental spray periods only adult surveys were carried out.

In the baseline pre-intervention circuit of each experiment (C1), study teams surveyed a single block together, proceeding as a group to an adjacent block until all houses in the study area were visited at least once. Houses that were not accessible on a day of a visit were revisited the next day and surveyed if open. After all study blocks were surveyed, houses that remained unsurveyed were visited a final time, and surveyed if possible. In subsequent circuits, similar spatially systematic surveying within subcircuits was carried out, and unsurveyed houses were visited a minimum of 3 times per circuit, or until access was obtained or refused.

### Experiment 1 (S-2013)

The initial S-2013 baseline pre-intervention circuit (C1) was carried out from 22-29 April 2013 in the spray sector, and from 29 April-16 May 2013 in the buffer sector (C1, Table 1, Fig. 3A). During the experimental treatment circuit (C2), Alphacypermetrin 10% (™Turbine 10%) was applied once per week for 6 consecutive weeks using Solo backpack sprayers (Cycles 1-6) or Colt hand-held sprayers (Cycles 4-6). Adult surveys were typically carried out during the spray period on Monday afternoons just prior to the initiation of each spray cycle, as described above. This design, therefore, measured adult densities up to 7-days after a previous spraying event. Post-intervention surveys (C3-C4) were initiated 10 days after completion of the last spray cycle (see Fig. S7A and S8A for detailed maps of surveys and sprays, respectively).

### Experiment 2 (L-2014)

Following the initial L-2014 baseline, pre-intervention circuit (C1), the experiment was interrupted by a MoH citywide emergency intervention in response to a dengue outbreak (see also Text S1). The MoH intervention consisted of 3 cycles of indoor cypermethrin 20% (®SERPA ciper 20 EW) spray applied between 04:00-09:00 or 17:00-20:00 with Solo backpack sprayers. MoH personnel generally sent an advance team with loudspeakers announcing the arrival of the spray teams, who visited each house on a block a single time. The MoH personnel had no mechanism to spray houses missed on their initial visit. In contrast to S-2013, during the L-2014 baseline circuit (C1) study teams worked in two groups (4 two-person teams). To survey both sectors simultaneously, one group was assigned to the spray sector, while the other was assigned to the buffer sector.

In response to information from the MoH about their imminent emergency spraying program (above), we adapted our study design in 3 ways (see also Text S2). First, we coordinated with the MoH to conduct adult surveys on a subset of L-2014 houses prior to (~20% of houses, C2) and during the emergency spray period (~20% of houses in each spray cycle, C3). No immature surveys were conducted during these circuits (for details see Fig. 3A, Fig. S7B, and Table S1). Second, we conducted independent monitoring of the 3 emergency citywide spray cycles (C3), along with standard quality control spraying procedures. We added a circuit of four spatially systematic subcircuits of full surveys (immature and adults, C4) during the MoH post-intervention period. Third, we added an extra circuit of adult surveys (~25% of houses, C5) that preceded experimental intervention. After Circuit 5, we resumed our planned L-2014 experiment (See Fig. S7B for a detailed map of survey locations).

As in S-2013, we applied 6 weekly cycles of ULV spraying (C6). A different pyrethroid insecticide, cypermethrin 20% (ESTOQUE^®^ 20 E.C., Tecnologia Quimica y Comercio S.A.) was used. For each cycle, spraying began on Monday evening using Solo backpack sprayers. We attempted to spray all accessible houses. Follow-up spraying of houses missed during the first day was carried out Tuesday-Friday between 07:00 and 20:00 using Colt hand-held sprayers (see also Text S2). In L-2014, adult surveys were typically carried out one day after a house was sprayed.

### Data Analysis

Unless otherwise noted, we analyzed only *Ae. aegypti* data, and used houses as the basic spatial units of observation. During experimental spray periods, we assigned a “spray status” indicator variable to each adult survey. “Prior spray” indicated that a spray application occurred in that house (prior to the survey) during the current or previous calendar week (otherwise, “no prior spray”). During L-2014, the relative timing between spray and survey was unclear for a limited number of surveys, which were designated as “timing unclear” (Tables S4 and S5).

### Statistical Models

For each experiment, a suite of statistical models was developed to estimate the impact of spray treatment on mosquito densities, proportion of infested houses, and population age structure (as determined from parity examination). With one exception, all comparisons and significance tests were conducted within-experiment.

We used two generalized linear model (GLM) specifications, both of which used a log link. For all counts, we used a negative binomial GLM (NB-GLM). Here, the response was the count of mosquitoes per house, and was assumed to follow a negative binomial distribution. The NB-GLM estimates the log of mean counts, and is akin to Poisson regression, while allowing for response over-dispersion (separate mean and variance) (Zeileis et al. 2008). For all proportions, we used a logistic GLM (L-GLM, i.e., logistic regression). Here, the response was the proportion of successes (out of total number of events), and was assumed to follow a binomial distribution. The choice of “success” was an arbitrary label applied to one of two mutually exclusive possibilities (presence or absence). The L-GLM estimates the log probability of success. For ease of interpretation, all model results were un-transformed after analysis and displayed in the original (unlogged) scale of observations.

To identify structural, pre-perturbation differences between sectors, we used an NB-GLM that estimated the number of *Ae. aegypti* adults per house (AA/HSE) in the baseline circuit (C1) in response to physical characteristics of houses, including building, floor, and roof construction, as well as number of containers, rooms, and surveyed rooms.

To assess the effect of spraying, we used an NB-GLM that estimated AA/HSE in response to circuit and spray sector. In addition, we used a companion L-GLM that predicted Adult House Index (AHI: proportion of houses with 1 or more *Ae. aegypti* adults) in response to circuit and spray sector. Finally, we tested the NB-GLM model formulation with alternate responses: female *Ae. aegypti* adults per house, and *non-Aedes* adults per house.

A NB-GLM was also used to estimate the effect of study year and spray status on AA/HSE. This model included only surveys conducted in the spray sector during experimental spray periods.

Counts from immature surveys and parity surveys were converted to proportions: container surveys yielded per-house proportion of positive containers (henceforth called the PrPC), which is also referred to as the container index. Parity surveys yielded the per-house proportion of nulliparous females (henceforth called the PrNF). Each proportional measure (PrPC, PrNF, and PrIH) was analyzed using a pair of L-GLM, weighted by the number of observations, with a separate model for each study year. Predictors included circuit and sector. The response was the log proportion of “successful” events per house, i.e., detection of positive containers or nulliparious females. The container model estimated the log proportion positive containers per house, log(PrPC), and the reproductive status model estimated log proportion nulliparous females per house, log(PrNF). We also model the total number of *Ae. aegypti* positive containers per house (PC/HSE) using an NB-GLM. Note that Breteau Index (BI) = 100*(PC/HSE).

To further evaluate the effect of spraying on mosquito densities, we employed contrast analysis (Lenth 2016) on the sector-by-circuit NB-GLM. We contrasted between circuits (spray sector only), and between sectors. The between-circuit contrast was complicated by temporal variation, either in extrinsic environmental factors, such as weather, or in intrinsic ecological processes, such as demographic stochasticity. The between-sector contrast was complicated by potential spatial ecological differences between sectors. More robust conclusions can be made if both types of contrasts provide similar assessments of the effect of spraying.

For the statistical models of adult, immature, and parity surveys, statistically indistinguishable groups and 95% confidence intervals (CI) of experimental group effects were estimated using least-squares means, also known as predicted marginal means, via the *lsmeans* R package (Lenth 2016). Tukey’s method was used to control the family-wise error rate (Lenth 2016).

### Human Use Statement

The study protocol was approved by the Naval Medical Research Unit Six (Protocol #NAMRU6.2013.0001) Institutional Review Board, which included Peruvian representation, in compliance with all US Federal and Peruvian regulations governing the protection of human subjects. IRB authorization agreements were established between the Naval Medical Research Unit Six and the University of California at Davis and North Carolina State University. The protocol was reviewed and approved by the Loreto Regional Health Department, which oversees health research in Iquitos. In all instances consent from adult members of houses was obtained without written consent. Written information sheets were provided to study participants, providing a detailed overview of the experiment design, procedures, and study goals before initial pre-interventions surveys. Permission to enter houses was provided at each survey or spray application visit.

## RESULTS

### Overview

In the six weekly ULV spray cycles of S-2013, 1,860 spray applications were carried out in 398 houses. During L-2014, 4,986 spray applications were carried out in 1,110 houses. A total of 3,843 surveys over 16 weeks and 12,124 surveys over 44 weeks were carried out in S-2013 and L-2014, respectively (Fig. 3A, Table 1). Adult *Ae. aegypti* densities were highly variable over space (Fig. S1) and time (Fig. S4) with highly skewed distributions. No adult mosquitoes were collected from most houses, and large numbers of adults were captured in very few houses (Fig. S1).

Model contrasts (AA/HSE) are shown in Fig. 5; details of adult densities and house indices are shown in Tables S6-S7. Overall, adult densities in the S-2013 baseline circuit (early May, C1) were 0.26 and 0.40 *Ae. aegypti* per house (AA/HSE) in the buffer and spray sectors respectively. During this same baseline circuit, 15% and 16% of houses contained one or more *Ae. aegypti* adults (AHI) in the buffer and spray sectors, respectively (Tables S7A-B). The L-2014 baseline circuit (January, C1) showed that *Ae. aegypti* adult densities were higher than in S-2013: 0.62 and 0.77 AA/HSE in the buffer and spray sectors, respectively. A later preintervention circuit in April (C5, prior to experimental spraying) yielded 0.44 and 0.67 AA/HSE in the buffer and spray sectors, respectively. The corresponding AHIs for these surveys were 31% and 34% in the spray and buffer sectors, respectively for January, C1, and 22% and 28% for April, C5.

**Figure 5. [contrast].**
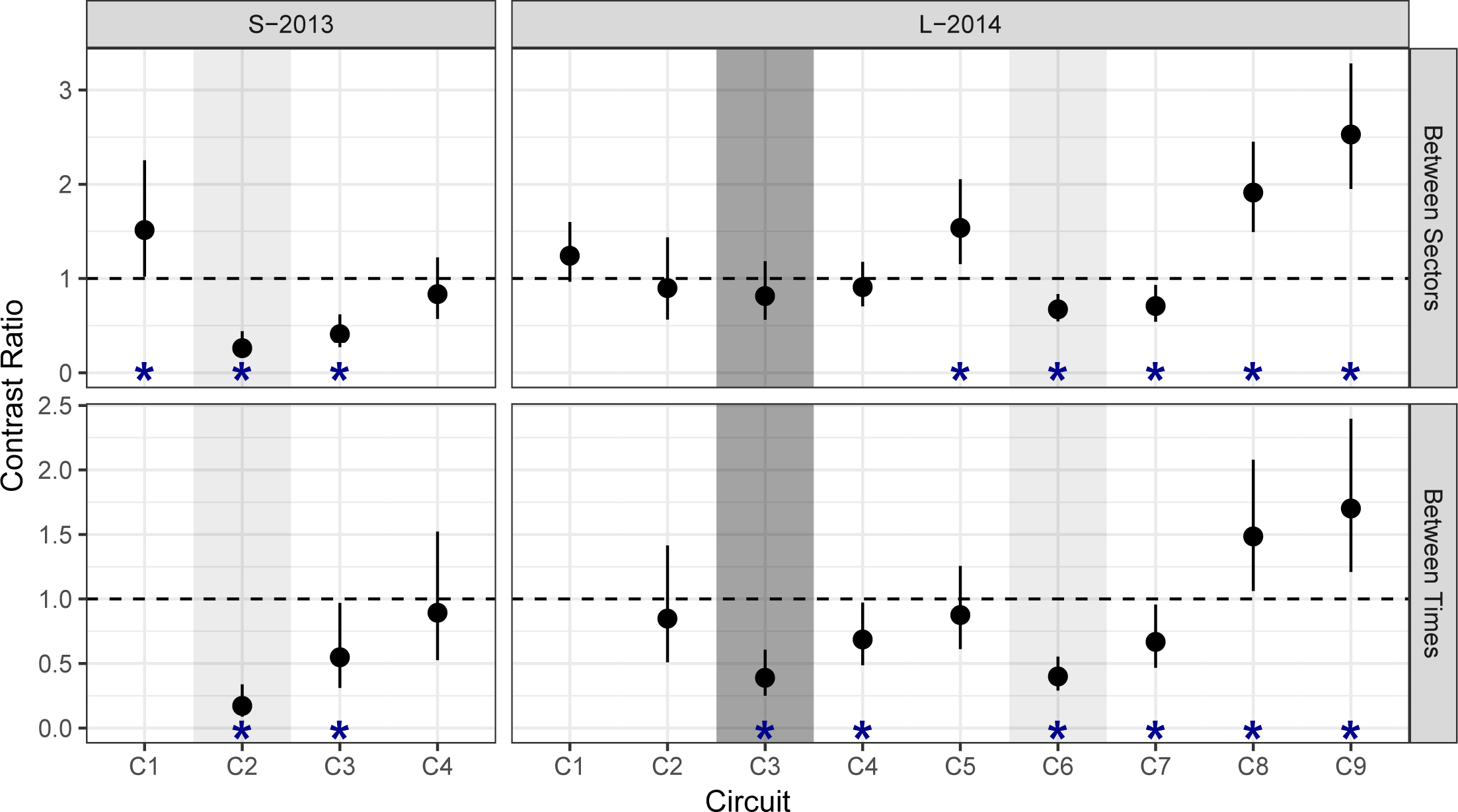
Contrast Ratios of AA/HSE, based on NB-GLM models shown in Fig. 4A. Top row (between-sector): Spray/Buffer. Bottom row (between-time, within spray sector): contrast relative to baseline (C1). Vertical bars show 95% CI. Horizontal dashed line indicates *H*_0_ of equality (ratio = 1). Asterisks (*) indicate significant difference between pairwise contrasts (reject *H*_0_).

Adult *Ae. aegypti* densities and house indices within the spray sector during spray periods were also lower during S-2013 (0.07 AA/HSE; AHI 5.5%) compared to L-2014 (emergency spraying, C3: 0.30 AA/HSE; AHI 18%; experimental spraying, C6: 0.31 AA/HSE; AHI 11%).

In the S-2013 post-intervention circuits (C3-C4), *Ae. aegypti* adult densities in the spray sector achieved a maximum of 0.35 AA/HSE (AHI 23%). In L-2014 (C7-C9), *Ae. aegypti* adult densities in the spray sector reached a maximum of 1.31 AA/HSE (AHI 41%).

### Meteorological conditions

Meteorological conditions were consistent between the two experiments, with average temperatures of 25.5°C (average minimum = 22.0°C, average maximum=32.0°C) and 25.6°C (average minimum = 22.0°C, average maximum=31.9°C) during the S-2013 and L-2014 experiments, respectively (National Climatic Data Center, https://www.ncdc.noaa.gov/cdo-web/). Precipitation during both years was approximately 0.84 cm per day. During the L-2014 entomological surveys for the MoH emergency citywide spray operation (January-March 2014), the temperatures were higher (average 25.9°C, average minimum = 23.3°C, average maximum=32.6°C) and it was rainier (average 1.09 cm per day) than at other times during the S-2013 and L-2014 experiments.

### Baseline Surveys

Comparisons of spray and buffer sectors in both experiments indicated that the two sectors had similar housing characteristics. No household physical characteristic was a predictor of adult mosquito density (data not shown). Consequently, we did not include such characteristics in our statistical models. Overall, for both years baseline numbers of *Ae. aegypti* adults were comparable between spray and buffer sectors (Table S2). During S-2013, however, we found a marginally significant difference between the buffer and spray sectors during the baseline (C1) circuit (0.26 vs 0.40 AA/HSE, resp.; Fig. 5, Table S2, p=0.039), making some statistical analyses of spray impacts conservative. During L-2014, baseline densities (C1) were not significantly different between the buffer sector (0.62 AA/HSE, AHI=31.1%) and spray sector (0.77 AA/HSE, AHI=33.7%) (Fig 5, Table S2, p=0.09). We observed no statistically significant baseline differences in adult female age structure between buffer and spray sectors (PrNF, Tables S8A and S8B). Baseline immature indices were similarly not different; for example, Breteau Indices (BI = 100 * PC/HSE) ranged from 9.4-10 in the buffer and spray sectors in both experiments (Table S9A and S9B). Container indices (i.e., percentage of water-holding containers infested with larvae or pupae, 100*PrPC) ranged from 3.9-4.1 in S-2013 and 3.1-3.3 in L-2014 (Tables S10A and S10B).

### Spray Coverage

The average percent of houses sprayed was lowest during the 3 MoH citywide emergency spray cycles in L-2014 ranging from 71% during cycle 1 to 62% in cycle 3 (Fig 3B). For S-2013, coverage started at 77% in cycle 1, decreased to 73% in cycle 3, and then improving in each subsequent cycle to 90% (cycle 6). For L-2014, coverage started at 74% in cycle 1, then modestly increased over time to approximately 82% in cycle 6 (Fig. 3B).

In both experiments, most spray sector houses were sprayed in more than 3 out of 6 spray cycles, and more than half of the houses were sprayed in all 6 spray cycles (Fig. S2). The primary reasons for not spraying a house were: house closed when personnel visited (3-16% for S-2013 spray, 19-28% for L-2014 MoH emergency spray, 7-16% for L-2014 spray), or residents did not allow access to the house (6-14% for S-2013 spray, 9-11% for emergency spray, 8-11% for L-2014 spray). During the S-2013 experiment, but not in L-2014, we recorded the reasons given by residents for refusing access. In many cases, teams were allowed access on subsequent visits. In early cycles, about one-third of the refusals cited a direct objection to fumigation, saying they did not believe it was effective or that the teams were not really using insecticide. In other cases, the reason given was inconvenience to the residents: eating, bathing, working, selling food, or that a sick person or newborn was in the house and could not leave. In some instances the homeowner was not present so consent could not be given.

### Spray Efficacy

During S-2013, 24-hour mortality of caged sentinel mosquitoes ranged from 87-97% with some variation across cycles (Fig. S3). Mean mortality was lower in L-2014, ranging from 53-87%. Overall, we observed a significant decrease in spray efficacy in L-2014 relative to S-2013 (Table 2). During S-2013, Colt hand-held ULV sprayers were used on 1/3^rd^ of the blocks during spray cycles 4-6. We observed higher mortality and knockdown in cycles 4-6, and less variation than was observed in cycles 1-3, which only included backpack sprayers.

**Table 2. [tab_cage].** Effect of year on *Ae. aegypti* control cage knockdown and mortality, showing a significant decrease in spray efficacy in L-2014. A separate logistic generalized linear mixed model (L-GLMM) was fit for each assay (separated by horizontal line). Year is a fixed effect. Spray cycle and house are nested random effects. Each cage contains 25 mosquitoes taken from a field-derived colony. See also Fig. S3.

| Experiment | Assay | nObs | Group | Est | SE | 95% CI |
| --- | --- | --- | --- | --- | --- | --- |
| S-2013 | Kill (24 Hours) | 112 | **a** | 0.94 | 0.01 | 0.90-0.96 |
| L-2014 | Kill (24 Hours) | 76 | **b** | 0.75 | 0.05 | 0.62-0.84 |
| S-2013 | Knockdown (1 Hour) | 112 | **a** | 0.94 | 0.02 | 0.86-0.97 |
| L-2014 | Knockdown (1 Hour) | 76 | **b** | 0.65 | 0.10 | 0.41-0.83 |

Droplet size (mean+SD) varied between experiments and sprayer type. Colt sprayers had smaller and more consistent droplets (19.1+12.6 μm) than backpack sprayers (29.2+19.5 μm). During the L-2014 MoH emergency spray, backpack sprayers were not properly calibrated, with an average droplet size of 39.8+25.8 μm. This improved to 20.6+14.1 μm in subsequent cycles. During the L-2014 6-cycle experiment, droplet size averaged 18.1+14.7 μm and 23.6+13.2 μm for Colt and backpack sprayers, respectively.

### Experiment 1 (S-2013)

Surveys conducted during the 6-week spray period (C2) generally occurred about one week after spraying. During the spray period, ULV spraying reduced adult *Ae. aegypti* population densities rapidly and significantly from 0.40 to 0.07 AA/HSE after six cycles of spraying (Fig. 4), yielding an 82.5% reduction relative to baseline (Fig. 5, Table S3, p<0.00001). The buffer sector, in contrast, had 0.26 AA/HSE both before (C1) and during (C2) the spray period.

**Figure 4. [hypoth_circuit2].**
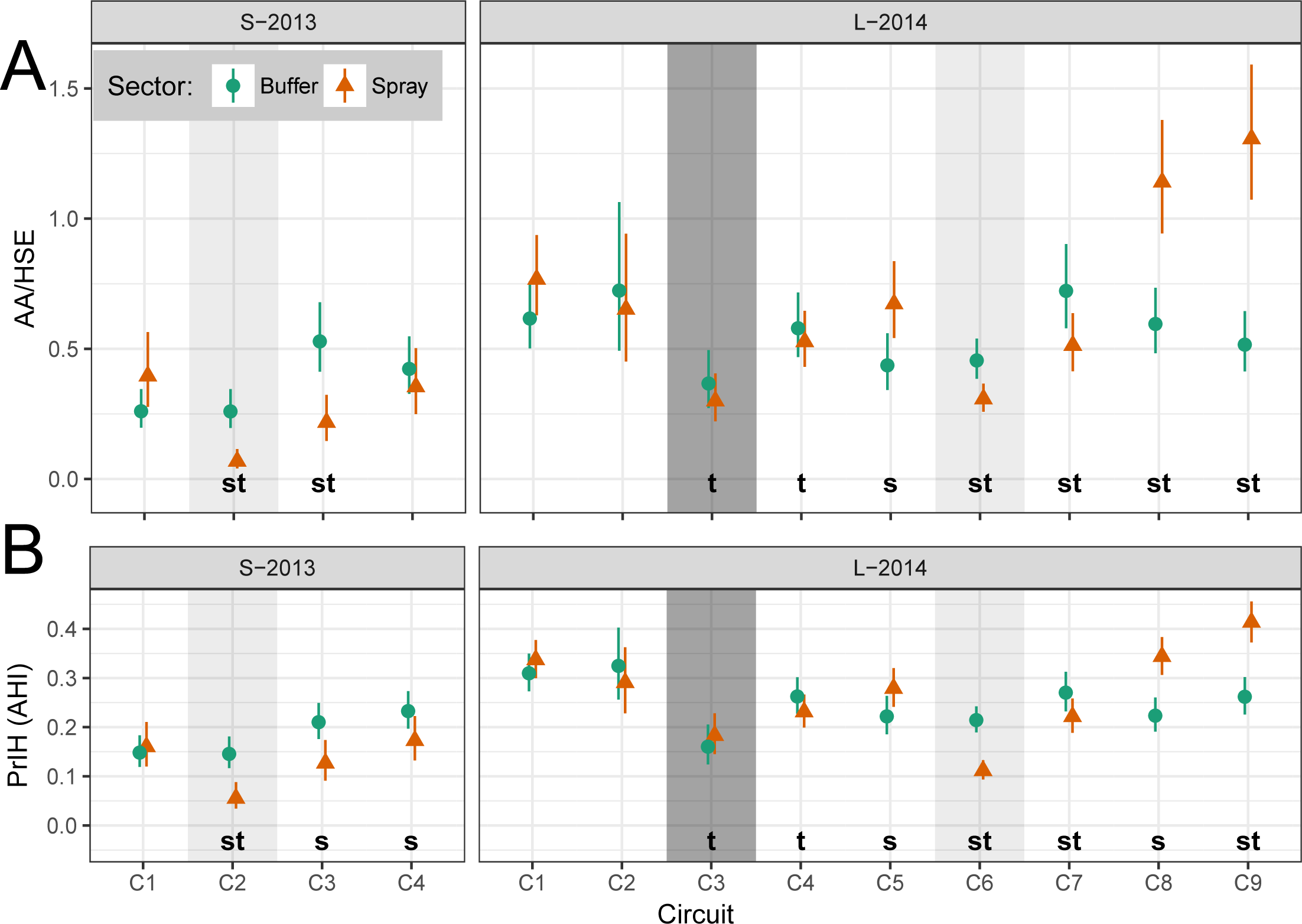
Model estimates of *Ae. aegypti* adults per house (AA/HSE, top row) and proportion infested houses (PrIH = AHI, bottom row). A separate generalized linear model (GLM) was constructed for each experiment (column) and for each measured response (row). **A: AA/HSE**: negative binomial GLM (NB-GLM). **B: PrIH**: logistic GLM (L-GLM). Models describe response of measure (row) to time period (X-axis) and treatment sector (color). Shading indicates spray events: experimental spraying (light) and citywide spraying (dark). Vertical bars show 95% Cl; non-overlapping Cl indicate highly significant difference. Letters (**s, t**) indicate significant differences between pairwise contrasts: **s**, between sector (within time, Table S2); **t**, between time (within spray sector, relative to baseline Cl, Table S3). See also Tables S6A-S6B, Tables S7A-S7B, and Fig. S5

Adult densities in the sprayed sector were 73.1% lower than in the buffer sector during the spray period (C2, Fig 5, Table S2, p<0.00001). Ongoing surveys within the spray sector during the spray period ranged from 0.04-0.08 AA/HSE, and did not change significantly over the course of the six sprays (Fig 6). Spray sector AA/HSE remained 45% lower than baseline levels during the first post-intervention period (C3, Fig. 5, Table S3, p=0.035), but densities increased from 0.04 to 0.27 AA/HSE between the first and second week post-spray. During the second post-intervention period (C4), spray sector adult densities returned close to baseline densities, increasing from 0.22 to 0.35 AA/HSE (Table S6A) which was 89% of baseline (Fig. 5, Table S3, p=0.94) and 83.3% of the buffer sector density at that time (Fig. 5, Table S2, p=0.36).

Adult house indices in the spray sector, by comparison, decreased from 16% during baseline surveys to 5.5% during the spray period (C2), then increased to 12.7% and 17.3% during the first and second post-intervention periods, respectively (C3-C4, Table S7A). In the buffer zone, AHIs were 15% during both baseline and spray periods, then increased to 21% and 23% in the first and second post-intervention evaluations (Table S7A).

During the S-2013 spray period (C2), only a small number of females (9 total) were collected in the spray sector (Table S8A). Therefore, we did not attempt to compare the age structure of Ae. *aegypti* populations before and after spray applications for this experiment. Model estimates of the proportion of nulliparous females (PrNF) showed accordingly high uncertainty (Fig. S5B and Table S8A).

Results from pupal demographic surveys followed a pattern similar to that of adult house indices. Baseline BIs were 10.0 in both the buffer and control sectors (Table S9A). BIs were not measured during the spray period, but during the first post-intervention period (C3) BI decreased slightly in the spray sector to 7.4 and increased to 16.1 in the buffer sector. During the second post-intervention period (C4), BIs were 15.1 and 22.3 in the buffer and spray sector, respectively. The post-treatment spray sector had statistically significantly higher PrPC than any other sector or time period (Fig. S5B, Table S10A). In the C4 spray sector, PrPC reached approximately 0.11, significantly higher than seen in the baseline spray sector (0.04) or in the C4 buffer sector (0.06).

### Experiment 2 (L-2014)

#### MoH Emergency Spray

MoH ULV spray applications were carried out in both experimental sectors (spray, buffer) prior to initiation of L-2014 experimental studies. In the baseline circuit (C1), AA/HSE ranged from 0.62-0.77 (Table S6B), and AHI ranged from 31-34% (Table S7B). During the citywide emergency spray period, AA/HSE decreased to 0.37 (AHI 16%) in the buffer sector and 0.30 AA/HSE (AHI 18%) in the spray sector, thus showing a modest 40-50% reduction in adult densities relative to the baseline Circuit 1 (Fig. 5, Table S3, p<0.0001). *Ae. aegypti* densities in the geographically central spray sector were more variable than for houses in the surrounding buffer sector. *Ae. aegypti* densities did show some recovery in the post-emergency circuit (C4), rising from 0.37 to 0.58 AA/HSE in the buffer sector and from 0.30 to 0.53 AA/HSE in the spray sector. There was also a small trend toward an increase in the proportion of nulliparous females (PrNF) between baseline and the emergency spray period, from 0.03 to 0.10 and from 0.07 to 0.11 in the buffer and spray sectors, respectively (C1 to C3, Table S8B).

Immature indices, which were measured at baseline (C1) and the post-emergency survey (C4), were similar over time. For example, the spray sector BI (Table S9B) during baseline (10.0) was not statistically different than in post-intervention surveys (6.3-11.9). The proportion of positive containers (PrPc) ranged from 0.4-0.5 across the baseline and post-emergency circuits (C1 and C4, Table S10B).

#### Experimental Spray

For our experimental evaluation, we carried out a circuit of pre-intervention adult surveys during April (C5) before initiating 6 cycles of ULV spray applications. In both spray and buffer sectors, adult densities were consistent with the January baseline surveys (Fig. 5, Table S3 p=0.95). During C5, however, there were significantly higher adult densities in the spray sector (0.67 AA/HSE) relative to the buffer sector (0.44 AA/HSE) (Fig. 5, Table S2, p=0.0034). During the experimental spray period (C6), AHI decreased significantly from 28 to 11% (0.67 to 0.31 AA/HSE) in the spray sector compared to the range of 22% and 21% (0.44 to 0.46 AA/HSE) in the unsprayed buffer sector (Table S7B).

Adult densities rebounded quickly after cessation of spraying (C7, Fig. 4, Fig. 6, Table S6B). AA/HSE increased from 0.31 during the spray period (C6) to 0.51 post-spray (C7), which was not statistically significantly different from the January baseline of 0.77 (C1) or from that of the April pre-intervention survey (C5, 0.67). During the L-2014 post-spray monitoring period (C7-C9), increases in adult densities were observed in the spray sector, with a 170% increase above January (C1) baseline levels in the final circuit (C9, Table S3). In the buffer sector, from C6 to C9, AHI ranged from a low of 21% during the spray period (C6) to a high of 27% (C7). In contrast, in the spray sector, AHI increased during each post-intervention survey, ranging from 11% during the spray period (C6) to 41% during the final post-intervention period (C9) (Table 7B). Adult densities during the first post-intervention circuit (C7) remained significantly lower than baseline (C1) levels (Fig. 5, Table S3, p=0.017). In C8-C9, however, densities were significantly higher than baseline levels (Table S3, p≤0.01). When comparing the buffer and spray sector, a similar pattern was observed. Adult densities during C7 remained significantly lower in the spray sector compared to the buffer sector. During C8 and C9, however, the spray sector had significantly more adult *Ae. aegypti* than the buffer sector (Fig. 5, Table S2).

**Figure 6. [boot_zoom].**
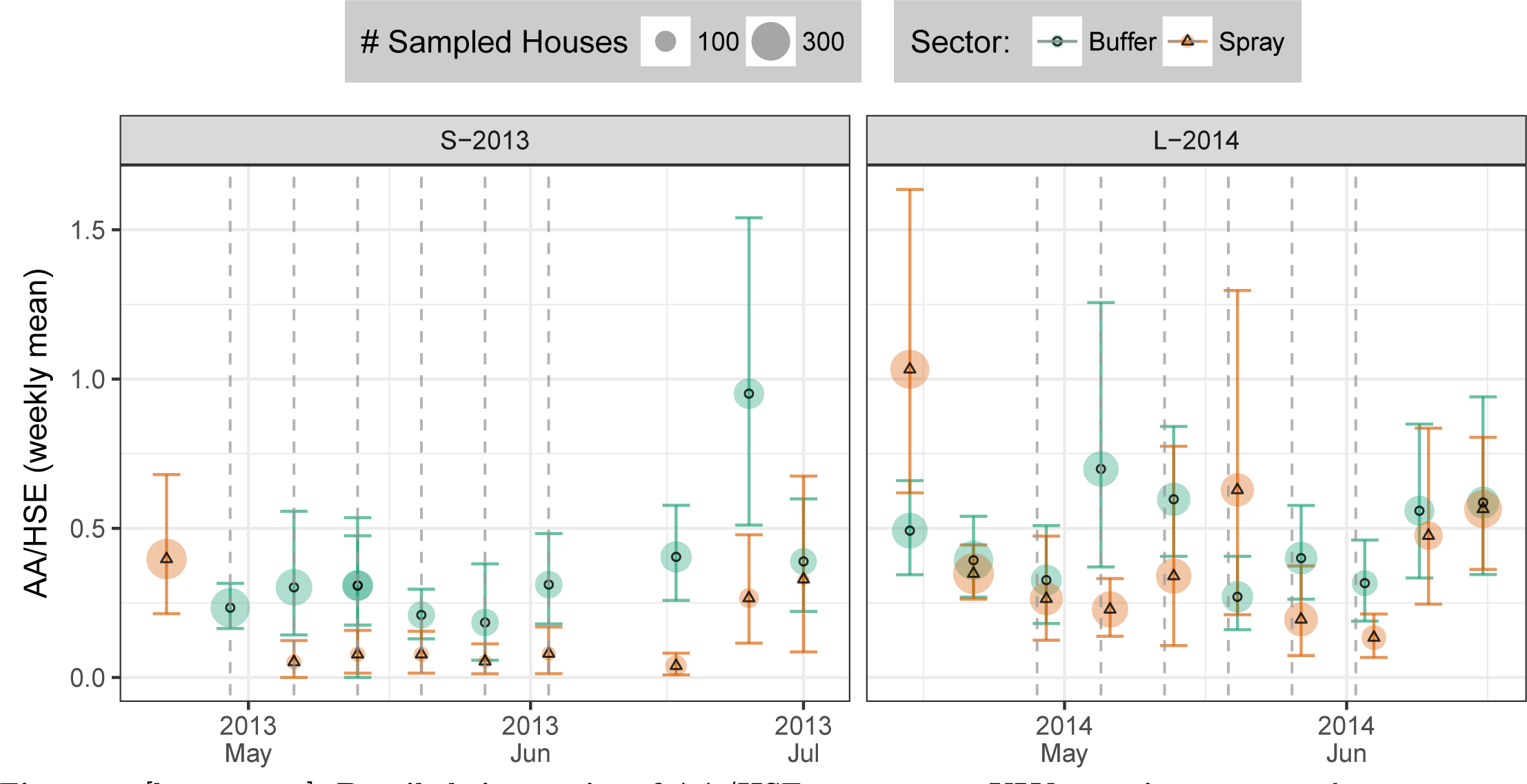
Detailed time series of AA/HSE response to ULV spraying, aggregated by week. X-axis shows week start date. Color and symbol shape shows sector (orange triangle: spray sector). Point size shows number of surveyed houses. Vertical dashed lines show approximate dates of experimental spraying (spray sector only). Vertical colored bars show bootstrap 95% CI (1e+04 draws per circuit).

Overall, we observed a strong effect of spraying on parity. During the spray period (C6), the proportion of youngest (nulliparous) females (PrNF) was significantly higher in the spray sector than in the buffer sector. Likewise, we observed an approximate doubling of PrNF in the spray sector relative to baseline (Table S8B).

Immature indices increased between the post-emergency spray survey (C4) and first post-experimental study survey (C7). For example, BI increased from 7.9 to 15.0 and from 8.3 to 11.9 in the buffer and spray sectors, respectively (Table S9B). Between the first and second post intervention surveys (C7-C8), BI dropped to 5.9 and 6.3 in the buffer and spray sectors, respectively. Two months later (C9), the BI decreased to 4.4 in the buffer sector and increased to 7.6 in the spray sector. Similar patterns were seen for the proportion of containers with immatures (Table S10B).

### Comparison of sprayed and unsprayed houses

During the S-2013 experiment, entomological surveys were carried out during the afternoon before each ULV spray cycle was initiated. For the majority of houses (254 of 398, 64%), therefore, *Ae. aegypti* densities were measured 7 days after the previous spraying. Only 2% of houses were sprayed fewer than 5 days earlier. In contrast, during the L-2014 experiment, entomological surveys were typically conducted the day after each spray cycle. This difference was the result of logistical concerns, as L-2014 involved many more houses. For 164 of the 1,259 house sprays (13%), the exact timing of each house spray was not available. In addition, some of the houses were sprayed later in the spray cycle (Table S5). The majority of L-2014 house surveys occurred within 2 days of spraying, and all houses were surveyed within 4 days of spraying. Thus, the average interval between house spray and survey was shorter than in S-2013. In the spray sector in both experiments, AA/HSE were lower in houses that had been sprayed the prior week compared to those that had not. In S-2013, AA/HSE was 0.06 and 0.11 in houses with prior spray and no prior spray, respectively, while L-2014 experienced 0.28 and 0.56 AA/HSE in houses with prior spray and no prior spray, respectively (Table S5). A (marginally) significant difference in AA/HSE between spray status groups was observed only during 2014 (Table S4, p=0.047).

## DISCUSSION

Despite the lack of a well-informed evidence base (Bowman et al. 2016), vector control of *Ae. aegypti* is often described as ineffective yet continues to be widely practiced by public health programs [6, 12, 26, 59, 60] (Simmons et al. 2012, Bowman et al. 2016, James et al. 2011, Reiter 2014, Andersson 2015, Bowman et al. 2016). Increasing attention has been given to integrated vector management, community involvement, and sustainability (Wilder-Smith et al. 2017). There is increasing recognition, however, that programs lacking interventions specifically directed at adult mosquitoes are insufficient for suppression of dengue and other *Aedes*-borne diseases [20, 22] (Morrison et al. 2008, Achee et al. 2015). A WHO dengue Scientific Working Group identified “analysis of the factors that contribute to the success or failures of national programs in the context of dengue surveillance and outbreak management”, including vector control as a priority topic for future research [61] (Runge-Ranzinger et al 2016).

Through two large-scale experimental studies and an assessment of a MoH emergency intervention campaign, our study evaluated an adulticiding strategy that is embedded in many national Aedes-transmitted virus control programs. We observed a clear *Ae. aegypti* population reduction during the extended period of repeated spray applications. These reductions were, however, not sustained after cessation of spraying.

Our study design could not logistically include randomized replicates [59, 63] (James et al. 2011, Reiner et al. 2016, Wilson et al. 2015) because we focused on monitoring spraying in large neighborhoods of houses. A review of previous *Ae. aegypti* space spray studies (Esu et al. 2010) shows that each replicate included 50 or fewer houses so that movement of adults from surrounding houses could have impacted results. In contrast, we monitored spraying in large numbers of houses; more than 1,100 houses (up to 2,100 houses) during the two experimental interventions, and a MoH citywide emergency spray program. Our experimental design reduced the potential impact of adults moving into the sprayed sector from unsprayed locations. In the citywide spraying, all areas of the city were expected to have about the same decrease in *Ae. aegypti* densities so adult movement should not have impacted the recovery at all. There is clearly a tradeoff between degree of replication possible and the size of experimental units.

In order to maintain study quality, our experimental interventions were supervised by trained entomologists. Our monitoring of the impacts of the L-2014 citywide emergency spraying provides a realistic and complimentary effectiveness assessment under practical, public health circumstances. It is also important to note that our study was primarily designed to provide data that could then be used to evaluate a computer simulation model (Magori et al) under extreme perturbation conditions, which was a major reason for evaluating a single centralized spray sector surrounded by a buffer sector.

The effectiveness of pyrethroid applications varied between years, but was similar between citywide emergency sprays and experimental sprays in 2014. Interestingly, in all experiments adult *Ae. aegypti* densities decreased significantly after the first cycle of spraying then fluctuated at relatively low levels during the remaining spray cycles; that is, additional cycles did not lower mosquito densities further. In all three interventions, adult populations partially recovered within 2 weeks of spray cessation. The pattern of rapid recovery of the *Ae. aegypti* population in our study is consistent with a number of previous reports (Esu et al. 2010) [5]. Studies by Perich et al. [23, 24] (2001, 2003) in Honduras and Costa Rica showed an approximately 90 percent reduction in adults one week after spraying, but the effect of the treatment was no longer significant after 6-7 weeks.

In the two experimental suppression trials we could not definitively determine if recovery of population densities was from adults migrating in from the surrounding buffer sector and/or from new adults emerging from development sites within the spray sector. However, in the emergency citywide spraying, the recovery was similar to that in the experimental trials. This suggests that movement of adults was not the key factor. Mosquito densities after the L-2014 experimental spray were monitored for a longer period of time: 23-weeks post-spray in L-2014 versus 9-weeks post-spray in S-2013. During L-2014, the density of adults in the spray sector increased to well above that in the buffer. In L-2014, ULV spraying resulted in a higher proportion of nulliparous females, indicating a shift to a younger adult female age distribution. This indicates that the spray sector continued to have active larval habitats that were producing new *Ae. aegypti* adults. In S-2013, for example, 22 *Ae. aegypti* positive containers were identified in a single house during a post intervention survey, whereas the baseline survey of that house revealed only three containers total, of which only one was positive. This kind of variation illustrates the stochastic and dynamic nature of *Ae. aegypti* larval habitats (LaCon et al. 2014, Getis et al. 2003). The dramatic L-2014 post-treatment increase cannot, however, be explained by an outlier in the form of a “superproductive” household (Morrison et al. 2014). One possibility is compensation by the immature population due to a reduction in larval population densities, which led to reduced density dependent competition within containers and increased survival to adult emergence. This kind of rebound effect merits further investigation.

In L-2014, both emergency and experimental spraying had significant, but lesser impact on the adult densities than in S-2013, even though L-2014 post-spray surveys were conducted (on average) fewer days after spray applications. The L-2014 24-hour mortality of caged sentinel mosquitoes was lower than in S-2013, something that could be due to characteristics of the different insecticide used, changes in pyrethroid resistance levels in Iquitos mosquito populations between S-2013 and L-2014, and/or differences in spray quality between the two experiments. By the end of 2014, significant pyrethroid resistance was detected in Iquitos (Palomino, INS report). Although we did not detect pyrethroid resistance before the S-2013 experiment, we do not have similar assay information from populations evaluated just prior to the L-2014 experiment. It is possible, therefore, that the lower efficacy observed in the L-2014 experiment was due in part to resistance in the local *Ae. aegypti* population. By 2015 the MoH had abandoned use of pyrethroid insecticides for indoor spraying and switched to malathion in an effort to improve efficacy.

A strong argument can be made that logistical challenges associated with application of ULV spray over a larger sector in the L-2014 experiment contributed to lower efficacy. First, Colt hand-held sprayers were only used in L-2014 when initially unsprayed houses were revisited, whereas in S-2013 they were used on at least 33% of the houses. Colt-sprayers had significantly better and consistent droplet sizes than backpack sprayers. The L-2014 experiment was a much larger effort with at least double the number of backpack machines and MoH fumigators participating, and droplets were only evaluated on a fraction of the machines used. In addition, during the L-2014 experiment coverage rates were lower overall.

Our results demonstrate that intensive, carefully administered space spraying can temporarily decrease the number and average age of female *Ae. aegypti* in houses. These results support smaller scale studies showing space spray induced reductions in *Ae. aegypti* density (Perich et al. 2000, 2001, 2003; Koenraadt et al. 2007). When, where, and how ULV mosquito control leads to meaningful reductions in disease remains a critical unanswered public health problem for policy makers. Computer simulation models have been employed to inform outcomes in limited situations, such as pathogen strain invasions [62] (e.g. Newton and Reiter 1992). Certain tentative recommendations, however, can be made based on existing data. Emergency indoor ULV spray interventions have the potential to mitigate *Ae. aegypti*-transmitted viruses, but coverage must be maximized with multiple spray cycles per house; i.e., at least 3 spray cycles based on our experience in Iquitos (Morrison and Scott, unpublished data). Officials should have no expectations of sustained reductions in mosquito densities and must recognize that these sprays only have the potential to mitigate the immediate impact of an arbovirus outbreak. Quality control of spraying efforts and insecticide resistance testing must be an integrated component of national programs. Although these are not new messages (WHO citations), our study adds new data to the vector control evidence base that we hope will better inform intervention programs and, thus, help refine policy for the application of space spray as a public health response to *Ae.* aegypti-transmitted viruses.

## AUTHOR SUMMARY

*Aedes aegypti* is a primary vector for medically important viruses that typically resides within houses. Indoor, ultra low volume (ULV) adulticide space spraying is considered to be more effective in controlling *Ae. aegypti* populations than outdoor spraying, and is widely used in tropical cities. Given the widespread use of indoor ULV spraying in emergencies by municipal control programs, the lack of large spatial scale evaluations is problematic. We conducted two large-scale experiments to evaluate indoor ULV pyrethroid spraying in the city of Iquitos, Peru in 2013 and 2014, and we also evaluated a municipal spraying effort. Our results demonstrate that densities of adults can be reduced by ULV spraying, but that adult densities in sprayed areas return to approximately pre-spray densities in less than a month. These findings agree with results from previous, smaller scale experiments, and confirm that ULV spraying should be viewed as having a short-term impact on *Ae. aegypti* populations. We provide extensive detail regarding our experimental design and data collection so that our results can assist in establishing best practices for future assessments of ULV spraying efforts, as well as aid in testing predictions of mathematical models of *Ae. aegypti* population dynamics.

---

**Supporting Information for *Efficacy of Aedes aegypti control by indoor Ultra Low Volume (ULV) spraying in Iquitos, Peru***

**Figure S1. [adult_baseline].**
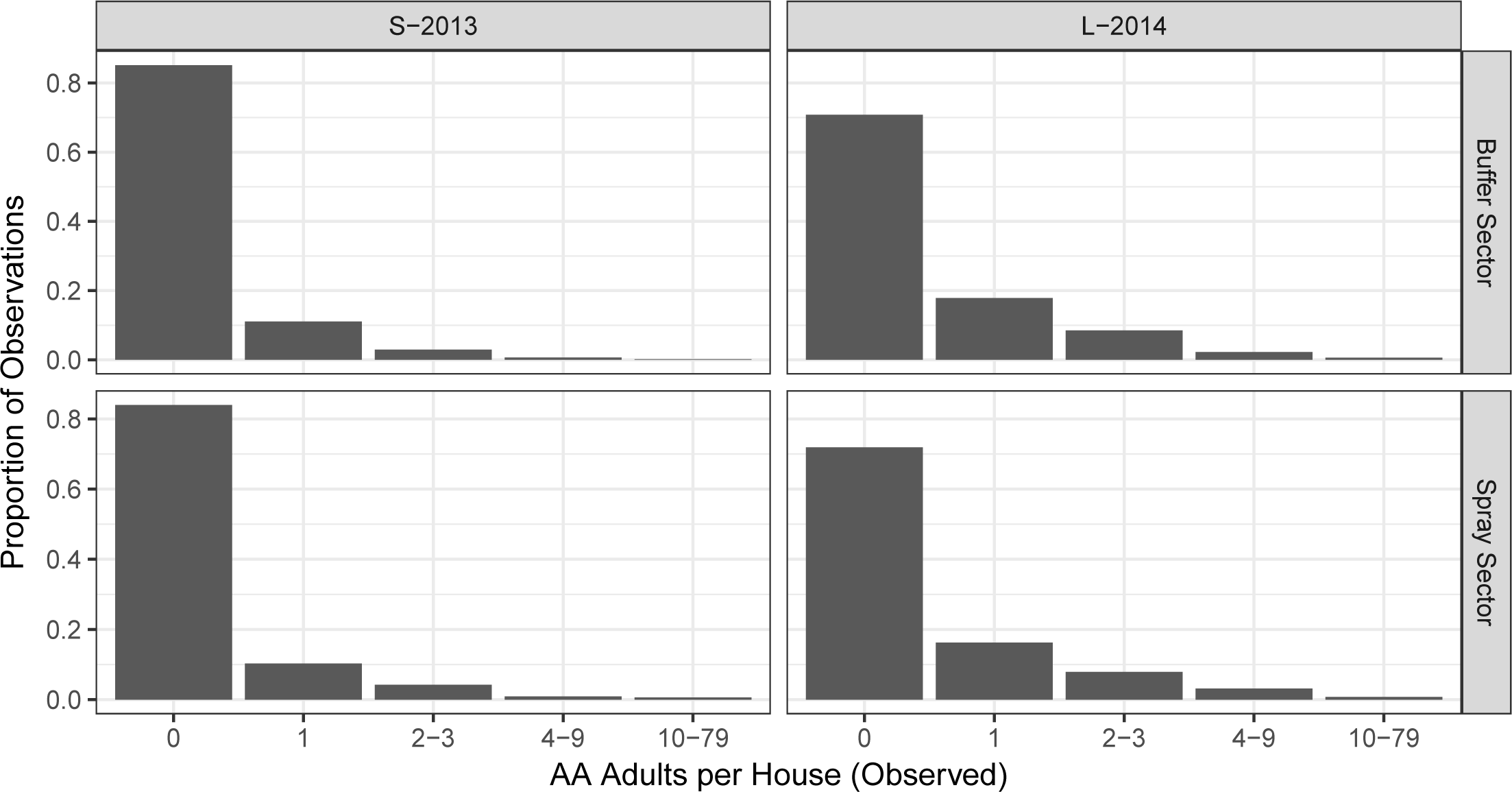
Histogram of AA/HSE at baseline (C1). Rows show treatment sector. X-axis is sqrt-scaled. The majority of house surveys find no adults.

**Figure S2. [spray_hist].**
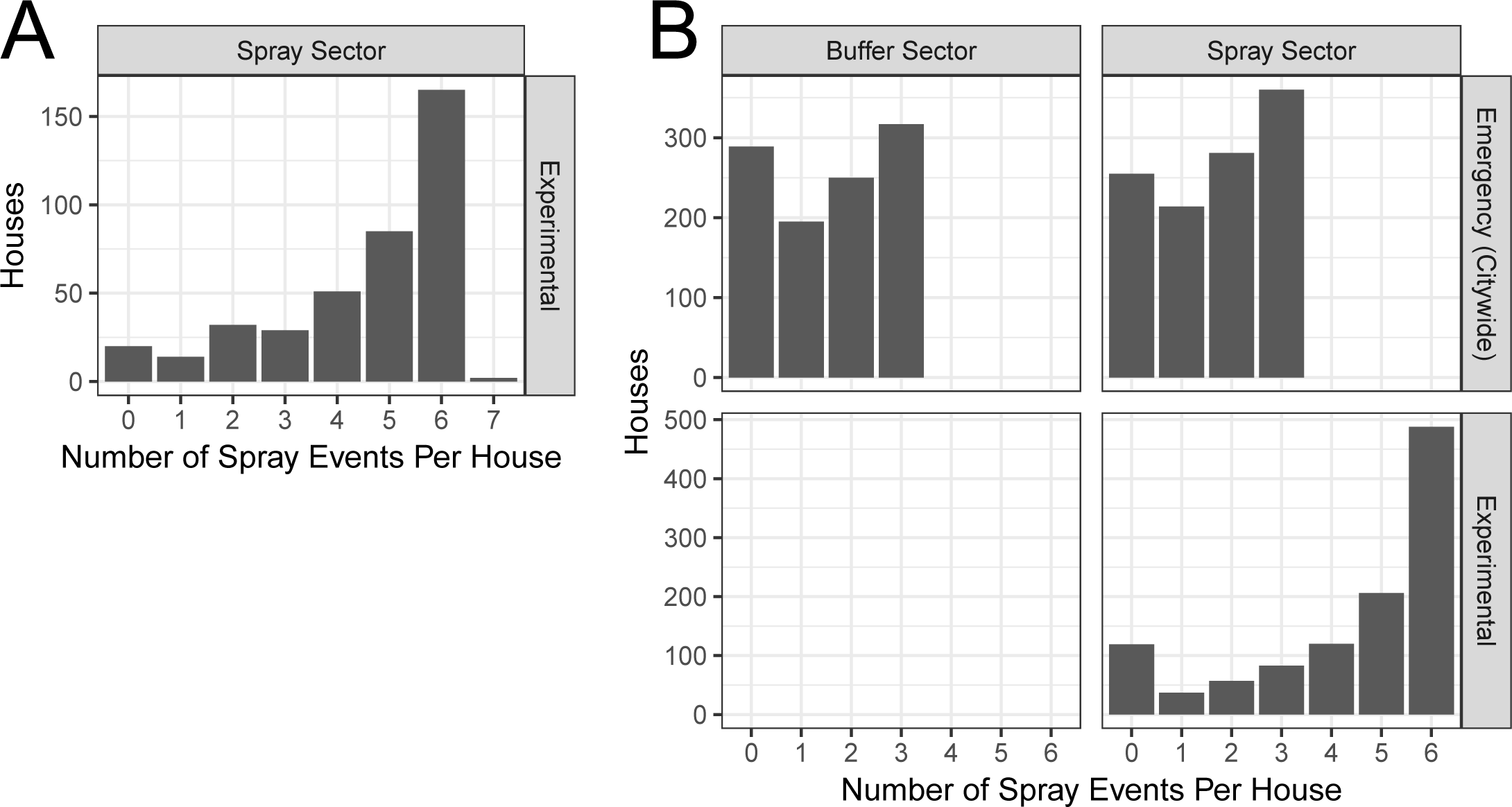
Summary of spray coverage in S-2013 (**A**) and L-2014 (**B**). In both years, most houses were sprayed in at least 5 out of 6 spray cycles, while a small number of houses were never sprayed. In L-2014, experimental spray coverage was much higher than emergency (citywide) spray coverage.

**Figure S3. [cage].**
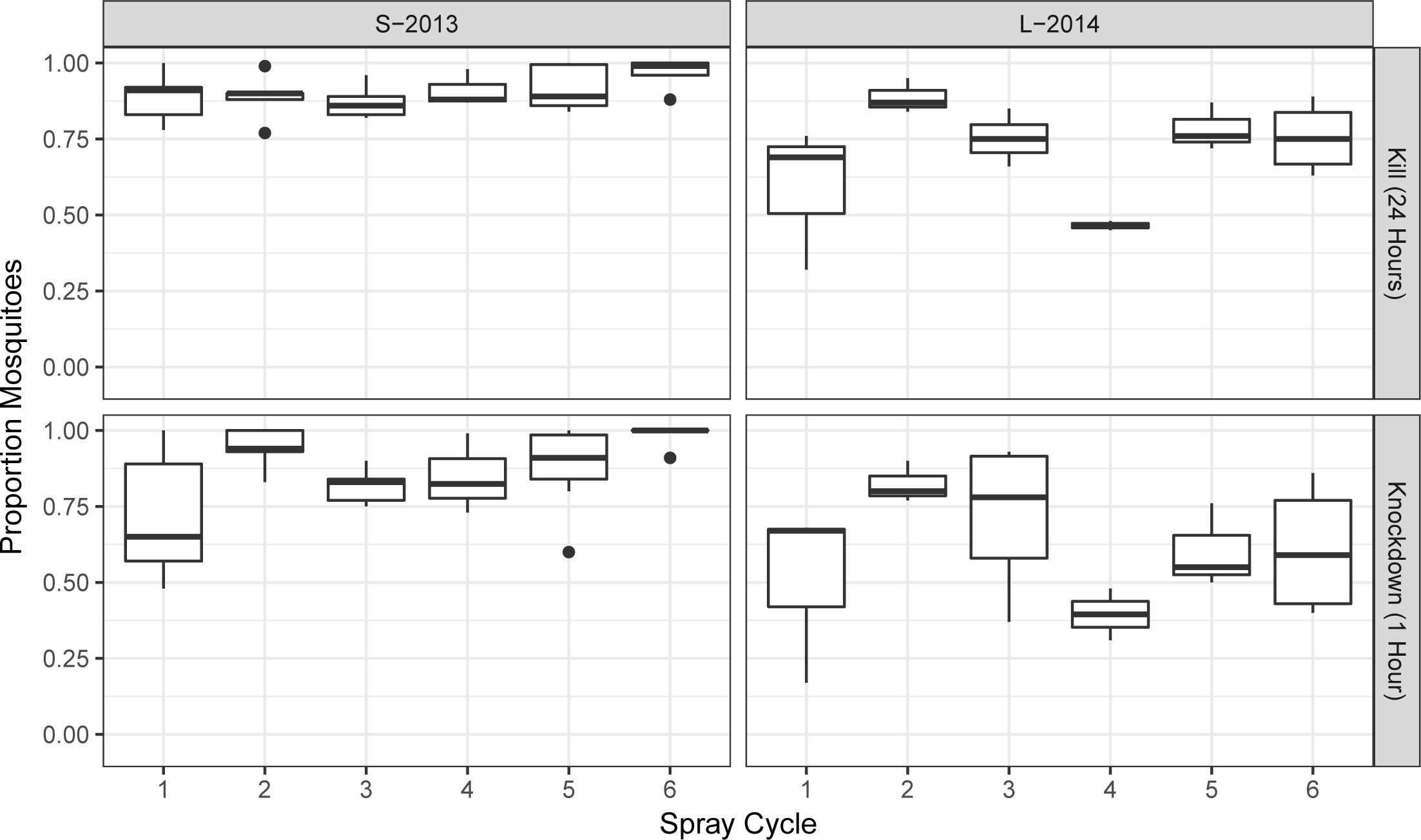
Boxplot of control cage house means: 25 adults per cage, 4 cages per house, approx 5 houses per spray cycle. Insects were from a laboratory colony (one colony per year).

**Figure S4. [ts].**
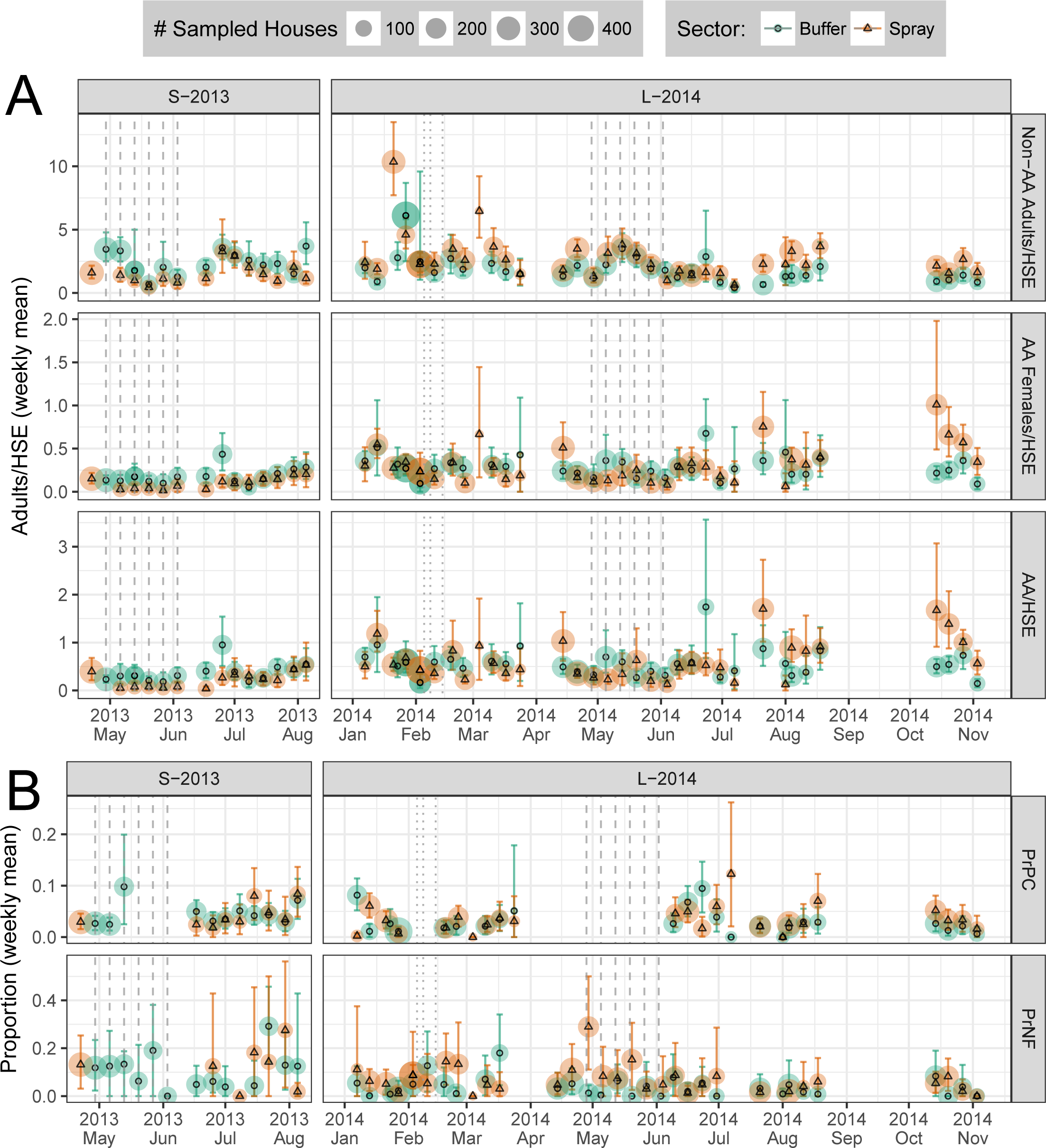
Time series of survey results, aggregated by week. X-axis shows week start date. Color and line-type shows treatment sector (orange dashed = Spray Sector). Point size shows number of surveyed houses. Vertical lines show approximate spray dates: dashed, experimental spraying (spray sector only); dotted, citywide spraying (Feb 2014, all sectors). Vertical colored bars show bootstrap 95% CI (1e+04 draws per circuit). **A**: Adult surveys. **B**: Container (PrPC) and Parity (PrNF) Surveys.

**Figure S5. [hypoth_full].**
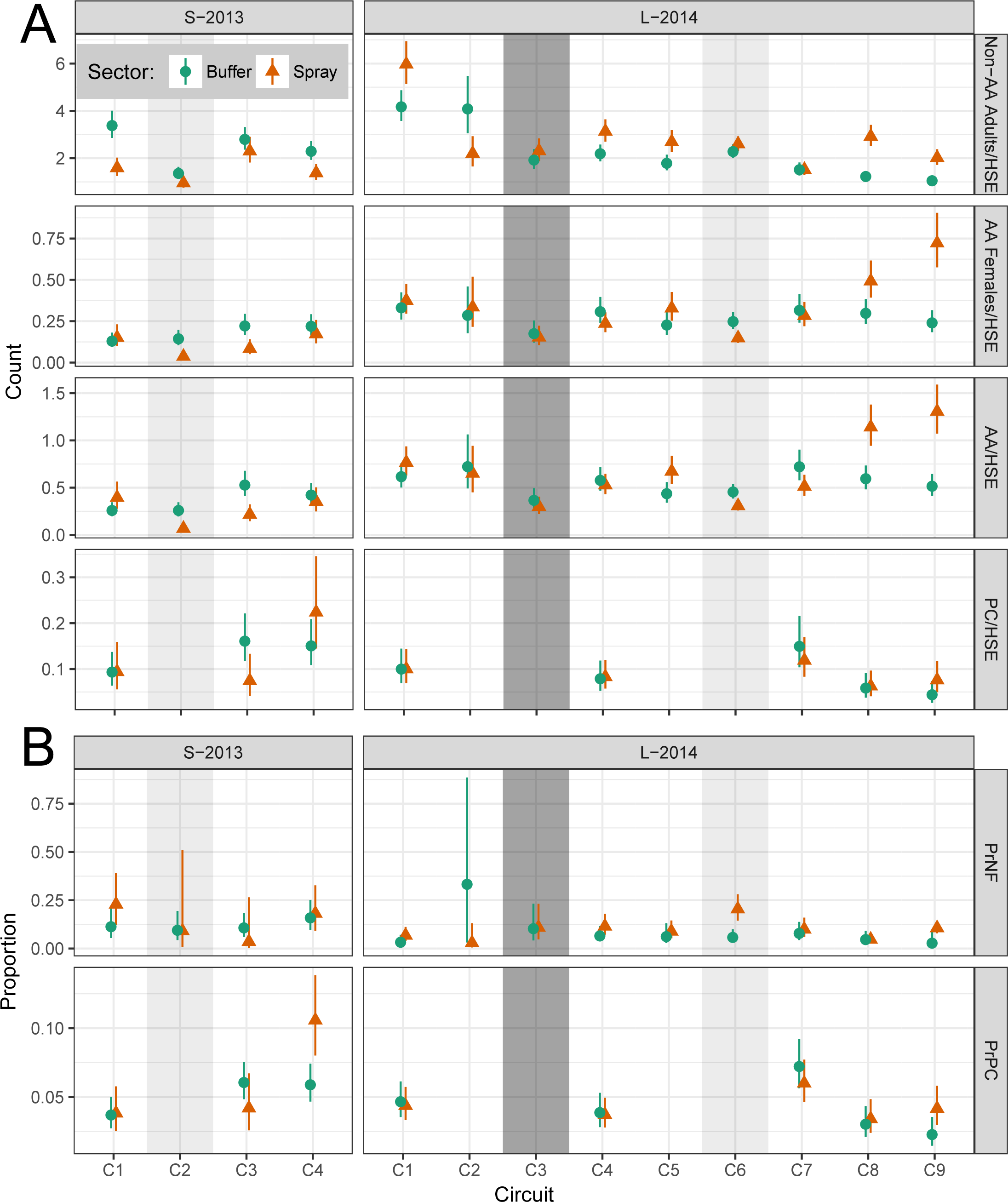
Model results, as in Fig. 4. All models include fixed effects of sector and circuit, with a separate model for each year. **A, Counts**: negative binomial GLM (NB-GLM). **B, Proportions**: logistic GLM (L-GLM). Note that Breteau Index (BI) = 100× PC/HSE. See also Tables S2-S10B.

**Figure S6. [map_base].**
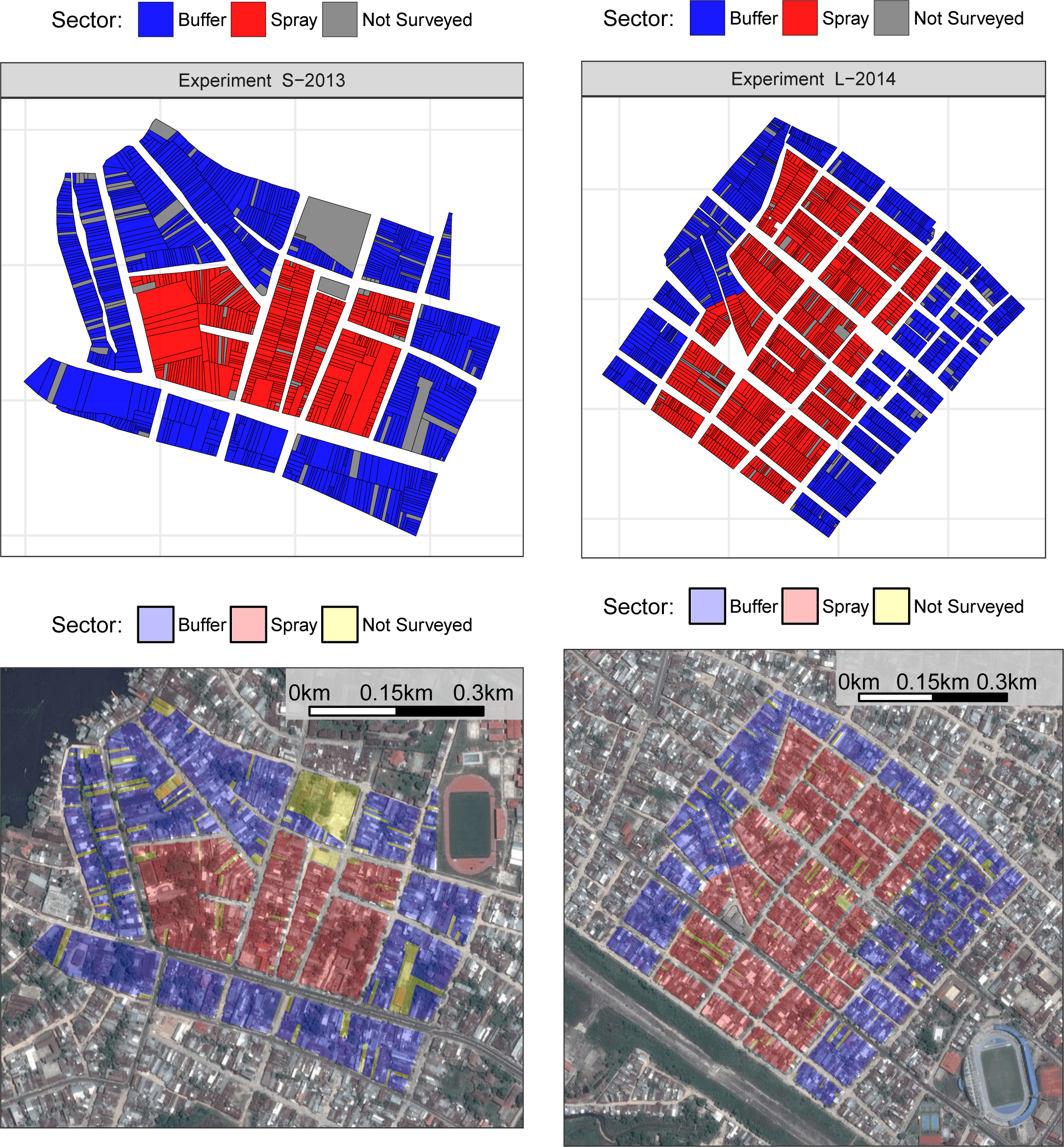
Maps of experimental areas, showing satellite imagery. Note the scale differs between experiments. See also Fig. 1.

**Figure S7. [map_week].**
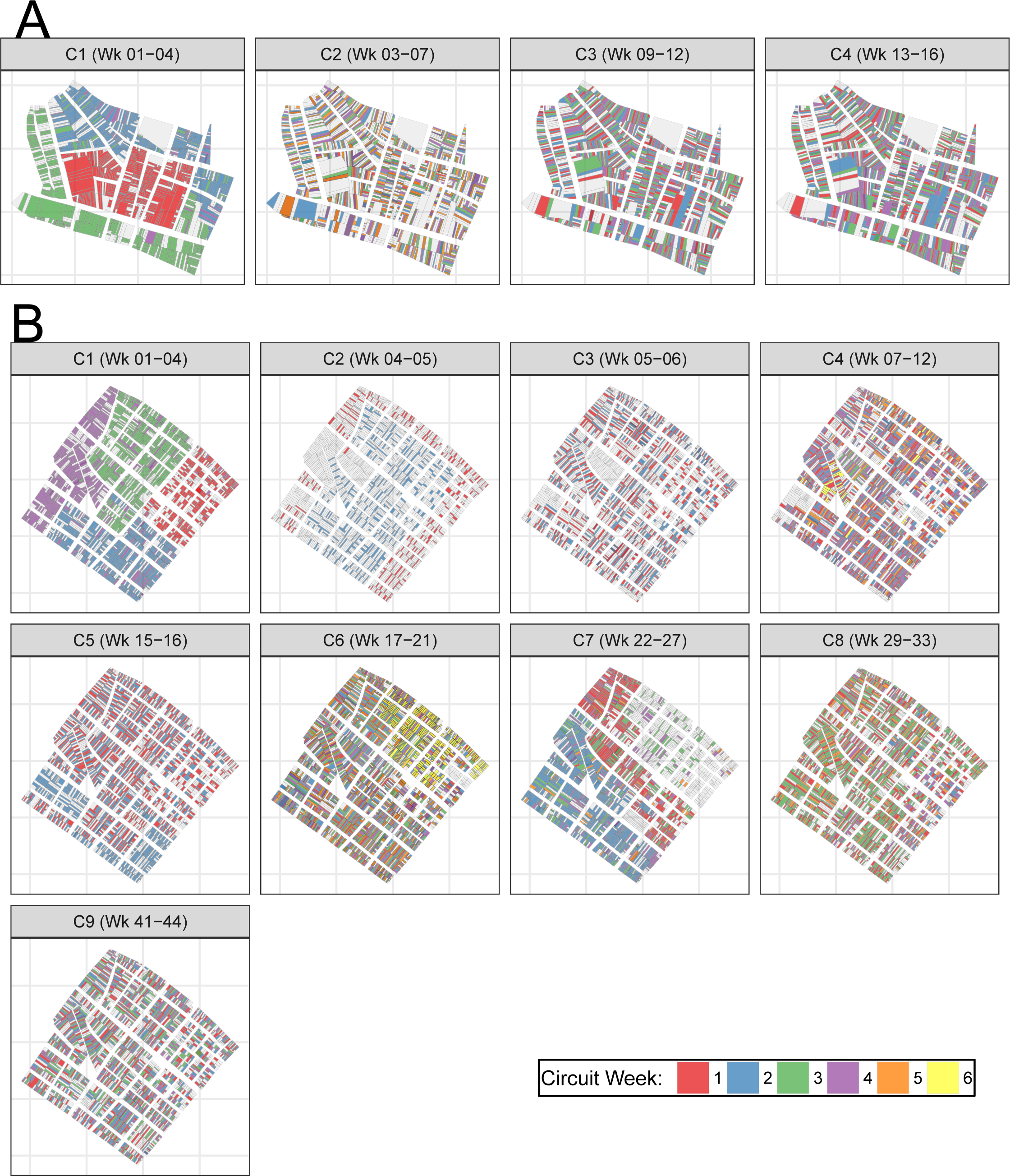
Map showing survey locations by circuit (panel) and week within circuit (color). **A**: S-2013. **B**: L-2014.

**Figure S8. [map_spray].**
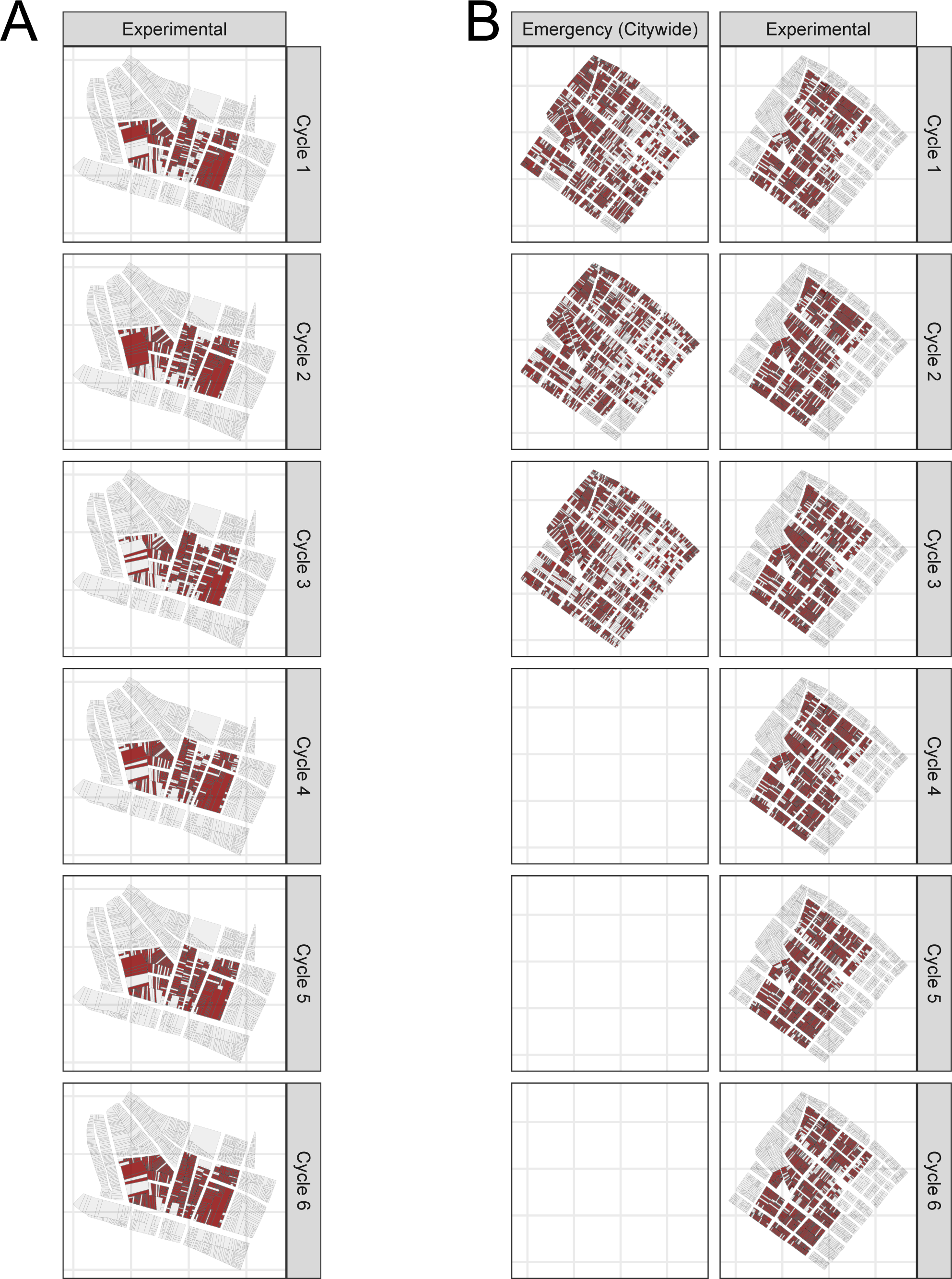
Maps of spray events (red) by spray cycle (rows). **A**: S-2013. **B**: During L-2014, 3 cycles of emergency citywide spraying were conducted, in addition to experimental spraying. Note the map scale differs between A and B. See also Fig. 1. S15

**Table S1. [tab_count_circuit2].**
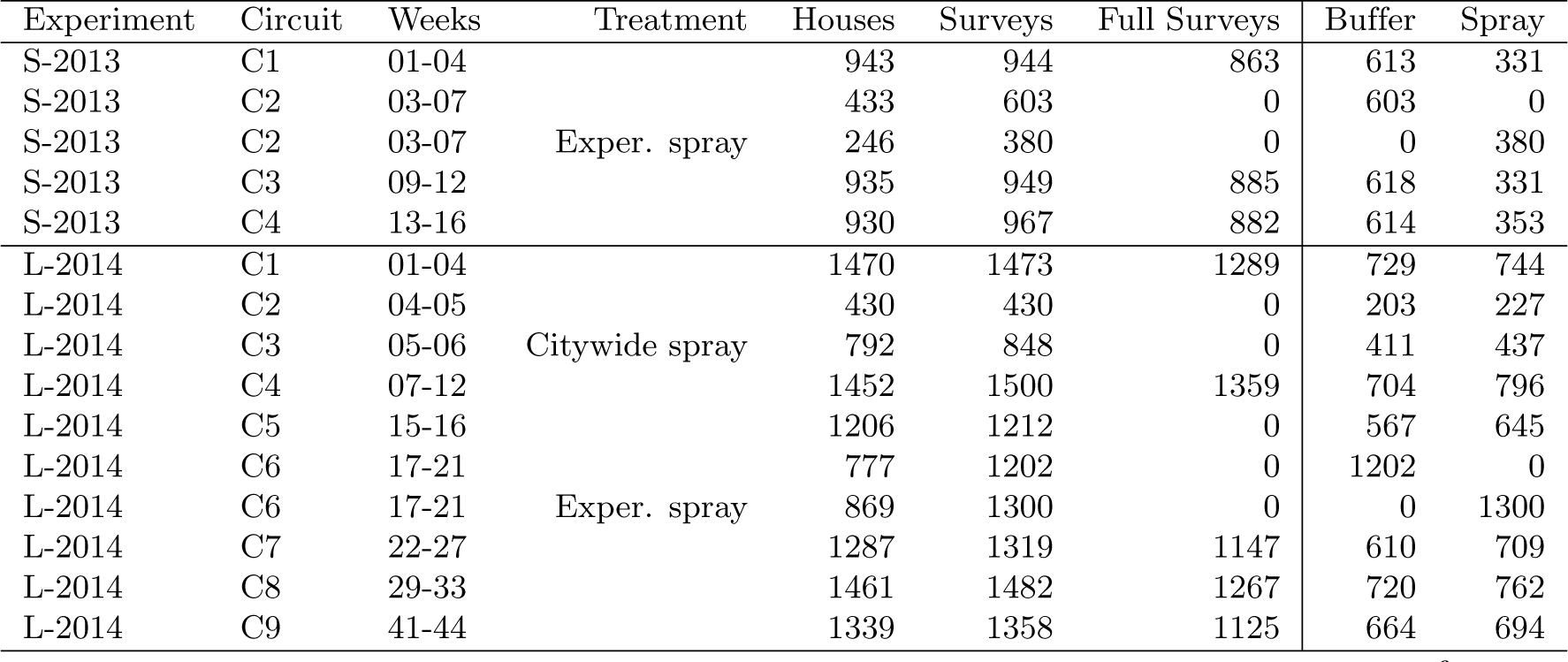
Observation counts by Circuit. *Weeks*: Week number from experiment start. *Houses*: number of unique houses surveyed. *Surveys*: total surveys (either adult, or combined adult and immature). *Full Surveys*: surveys where both adult and immatures were surveyed. *Buffer, Spray*: surveys in buffer and spray sector, respectively.

**Table S2. [tab_contr_tx].**
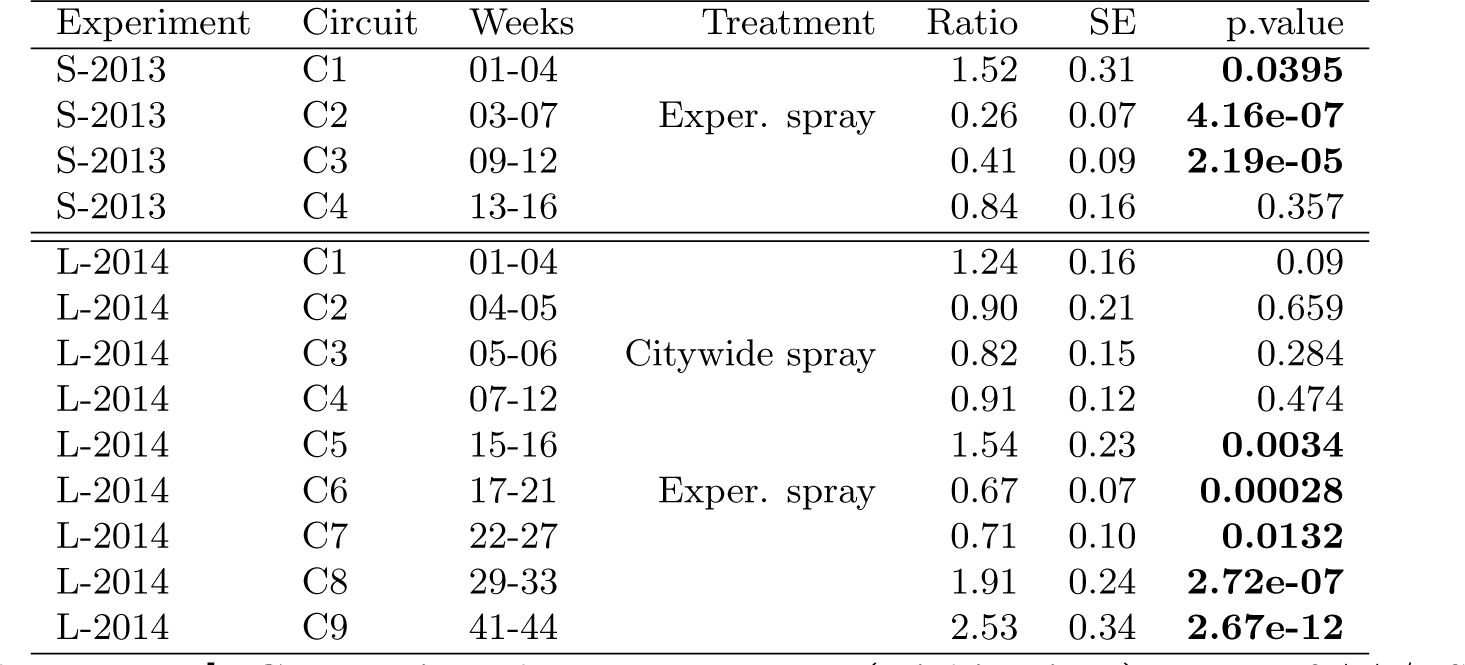
Comparison between sectors (within time): Ratio of AA/HSE in spray sector relative to buffer sector (spray/buffer). **Bold p.values**: significant difference between sectors. In both years, the spray sector starts with more adults per house, and spraying reduces AA/HSE relative to buffer sectors. As in Table S3, the effects of spraying are most pronounced in 2013. See also Fig. 4A.

**Table S3. [tab_contr_time].**
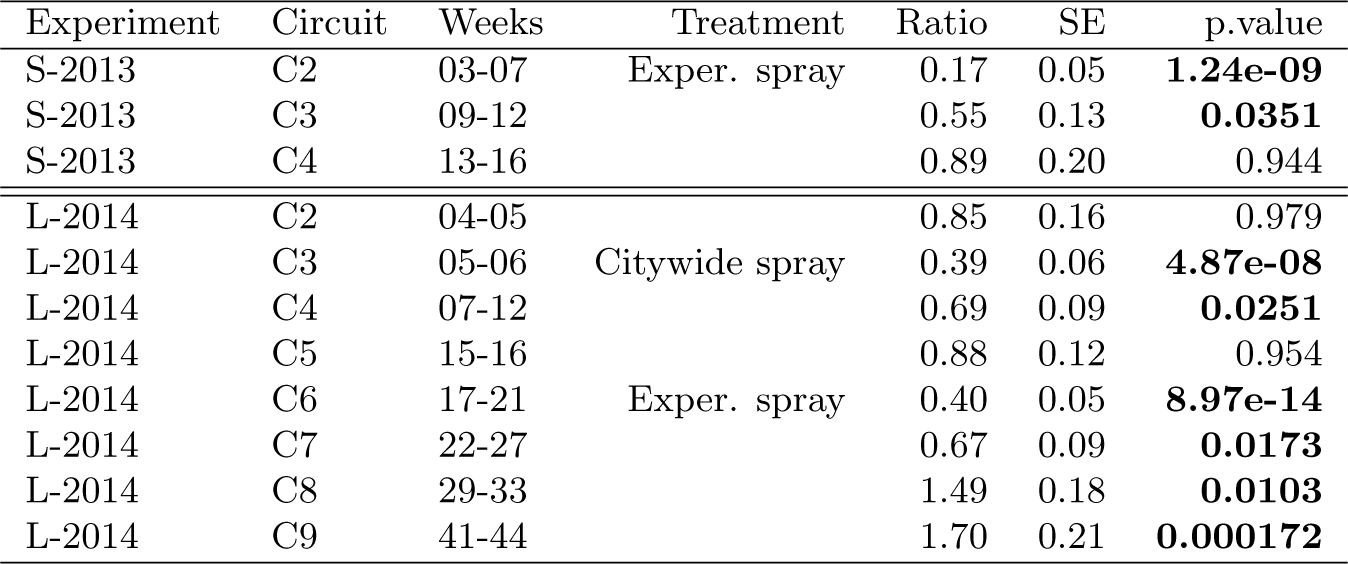
Comparison between times (within spray sector): Ratio of AA/HSE relative to baseline (C1, spray sector only). **Bold p.values**: significant difference from baseline circuit. In both years, spraying reduces AA/HSE relative to baseline (C1). The effects of spraying are most pronounced in 2013, but are short-lived in both years. See also Fig. 4A.

**Table S4. [tab_contr_spray].** Comparison between spray status (whether house was sprayed in prior week): Ratio of AA/HSE in houses that were or were not sprayed in the week prior to surveying (no prior spray / prior spray). **Bold p.values**: In L-2014, houses without prior spraying yielded significantly more adults than houses with prior spraying. In S-2013, most houses were sprayed in the prior week. In L-2014, the exact date of spraying was uncertain for a small number of houses. See also Table S5.

| Experiment | Contrast | Ratio | SE | p.value |
| --- | --- | --- | --- | --- |
| S-2013 | No Prior Spray / Prior Spray | 1.69 | 1.20 | 0.711 |
| L-2014 | No Prior Spray / Prior Spray | 2.01 | 0.62 | **0.0467** |
| L-2014 | Timing Unclear / Prior Spray | 0.71 | 0.26 | 0.568 |

**Table S5. [tab_hsd_spray].**
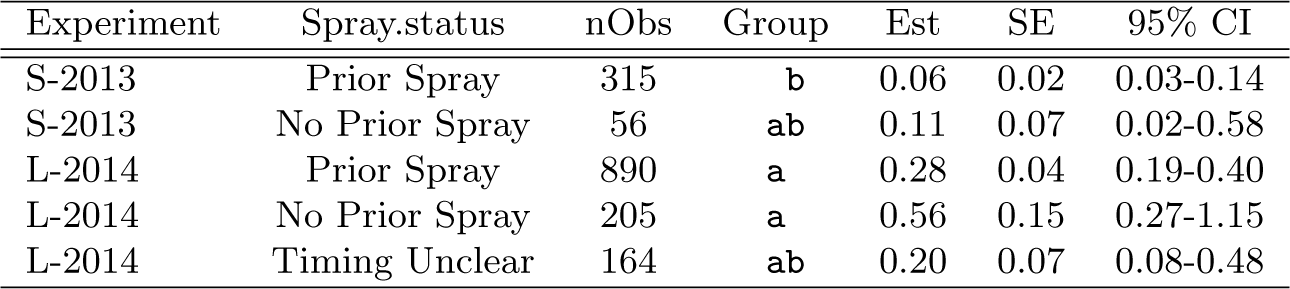
Effect of spray in previous week on AA/HSE. A single model (NB-GLM) includes both experiment year and spray status as predictors. Only house surveys in the spray sector during experimental spraying are included (i.e., S-2013 Circuit 2 and L-2014 Circuit 6). See also Table S4.

| Experiment | Spray.status | nObs | Group | Est | SE | 95% CI |
| --- | --- | --- | --- | --- | --- | --- |
| S-2013 | Prior Spray | 315 | **b** | 0.06 | 0.02 | 0.03-0.14 |
| S-2013 | No Prior Spray | 56 | **ab** | 0.11 | 0.07 | 0.02-0.58 |
| L-2014 | Prior Spray | 890 | **a** | 0.28 | 0.04 | 0.19-0.40 |
| L-2014 | No Prior Spray | 205 | **a** | 0.56 | 0.15 | 0.27-1.15 |
| L-2014 | Timing Unclear | 164 | **ab** | 0.20 | 0.07 | 0.08-0.48 |

**Table S6A. [tab_hsd_aedes_2013].**
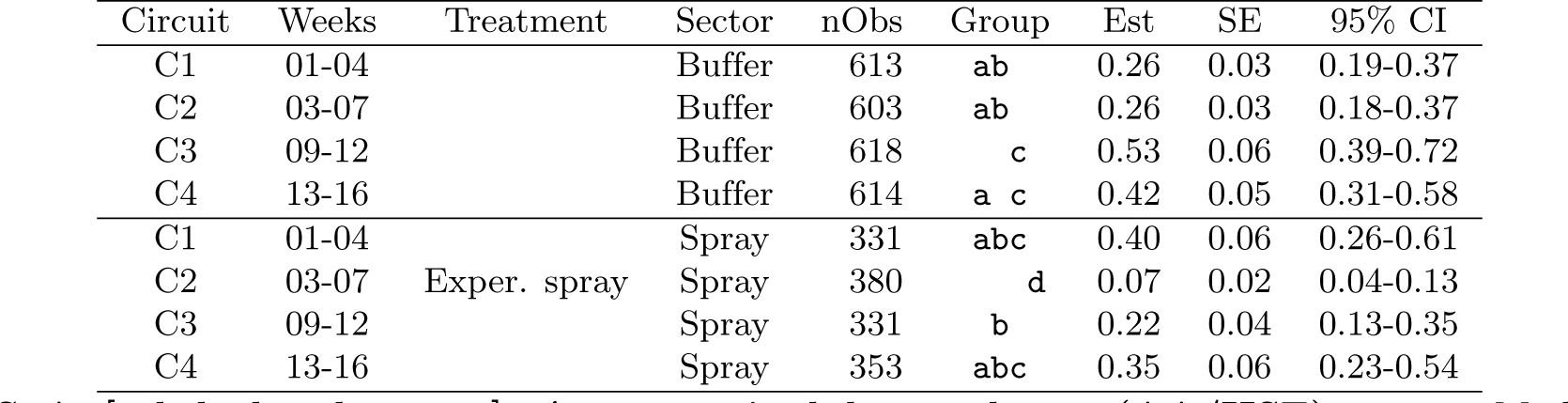
*Ae. aegypti* adults per house (AA/HSE), 2013. Model estimates by circuit and treatment sector. Horizontal line separates treatment sectors; significance groups (Tukey HSD) compare among all rows. See Fig. 4A for model description.

**Table S6B. [tab hsd aedes 2014].**
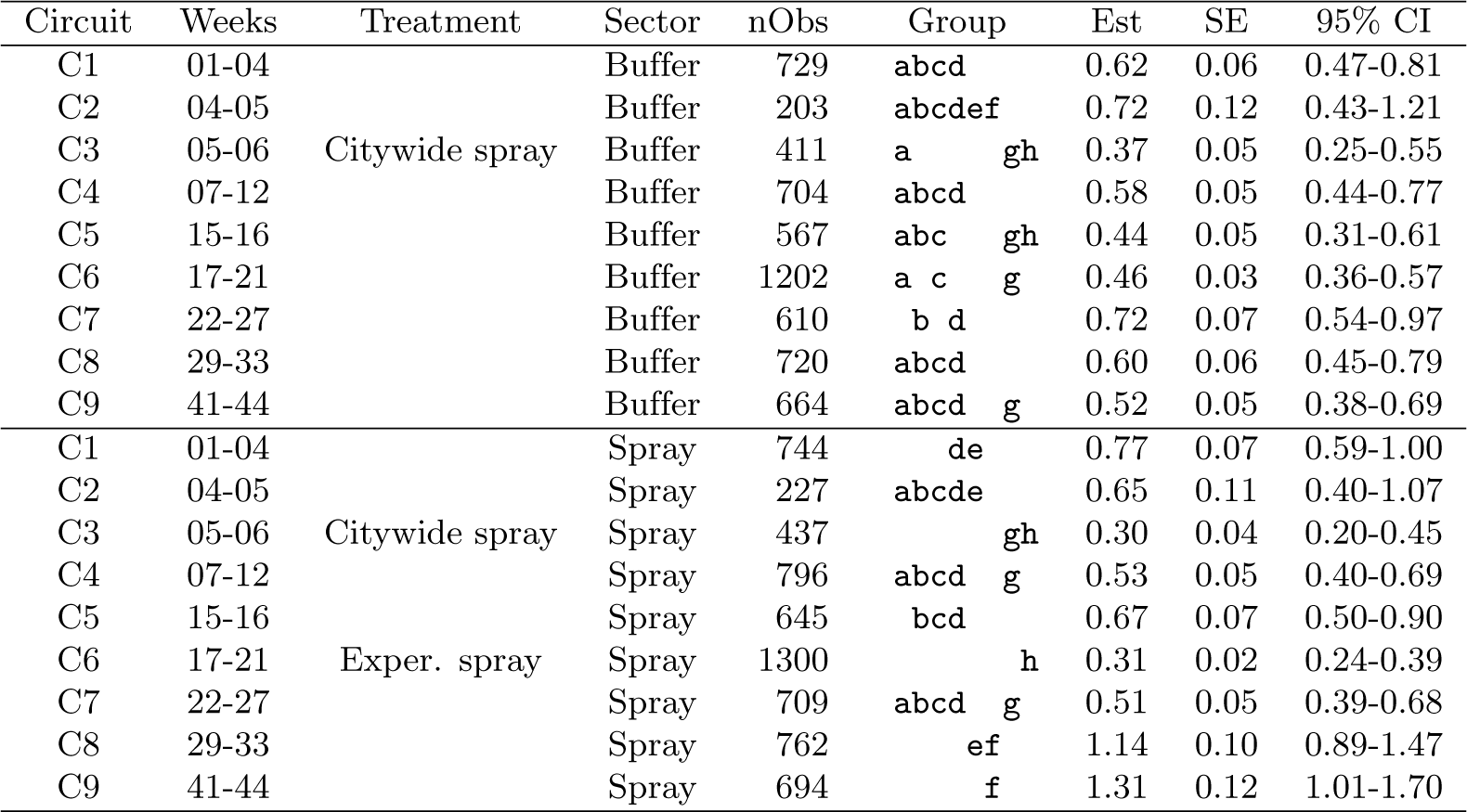
Ae. aegypti adults per house (AA/HSE), 2014. See Table S6A for details.

**Table S7A. [tab_hsd_infest_2013].**
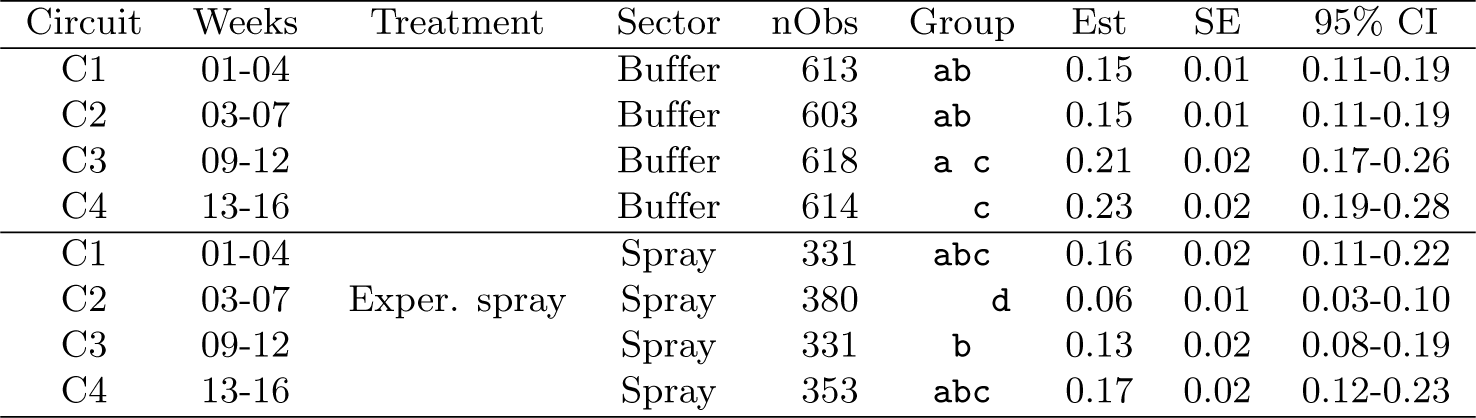
Proportion *Ae. aegypti* adult-infested houses (PrIH), 2013. Model estimates by circuit and treatment sector. Horizontal line separates treatment sectors; significance groups (Tukey HSD) compare among all rows. See Fig. 4B for model description.

**Table S7B. [tab hsd infest 2014].**
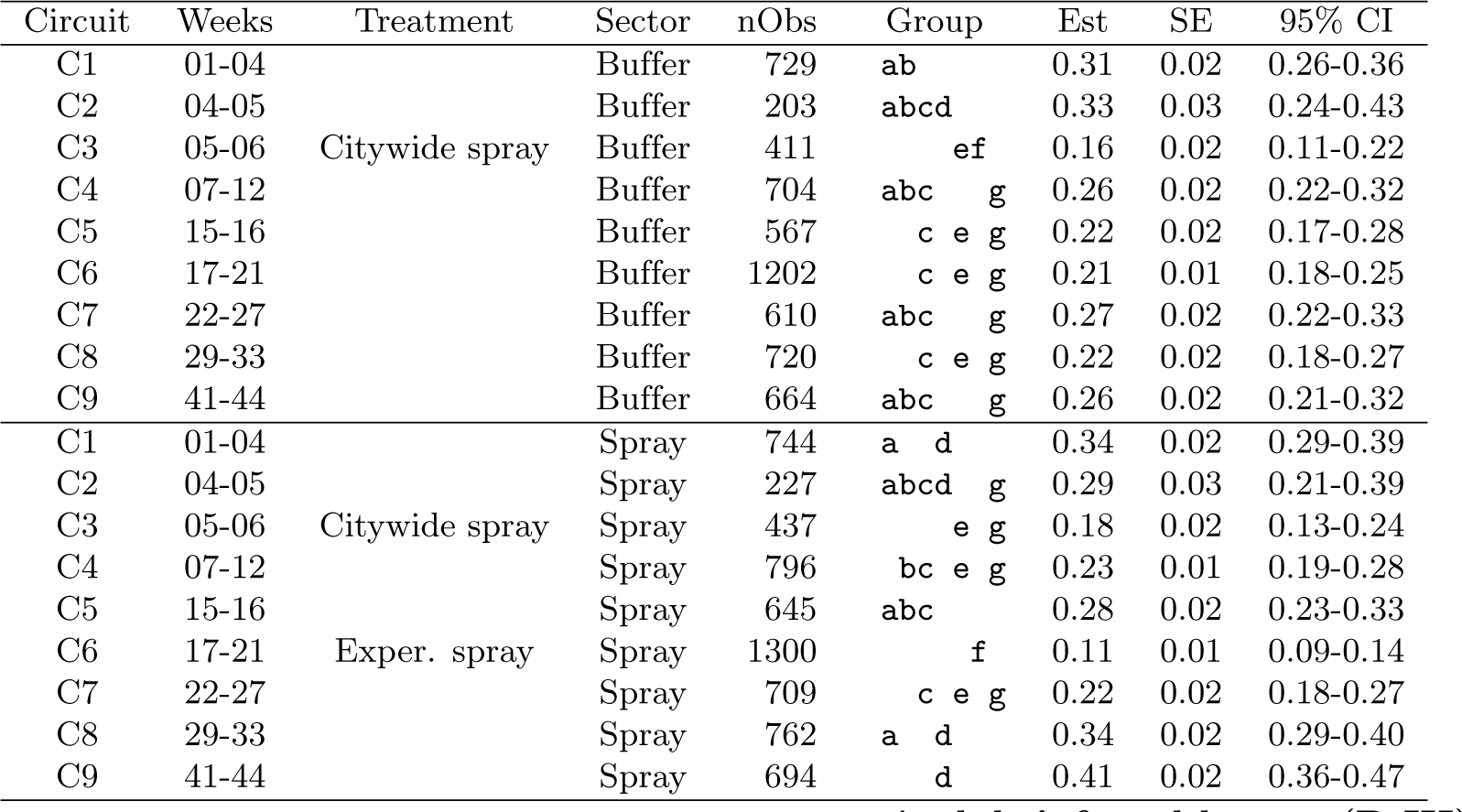
Proportion Ae. aegypti adult-infested houses (PrIH), 2014. See Table S7A for details.

**Table S8A. [tab_hsd_par_2013].**
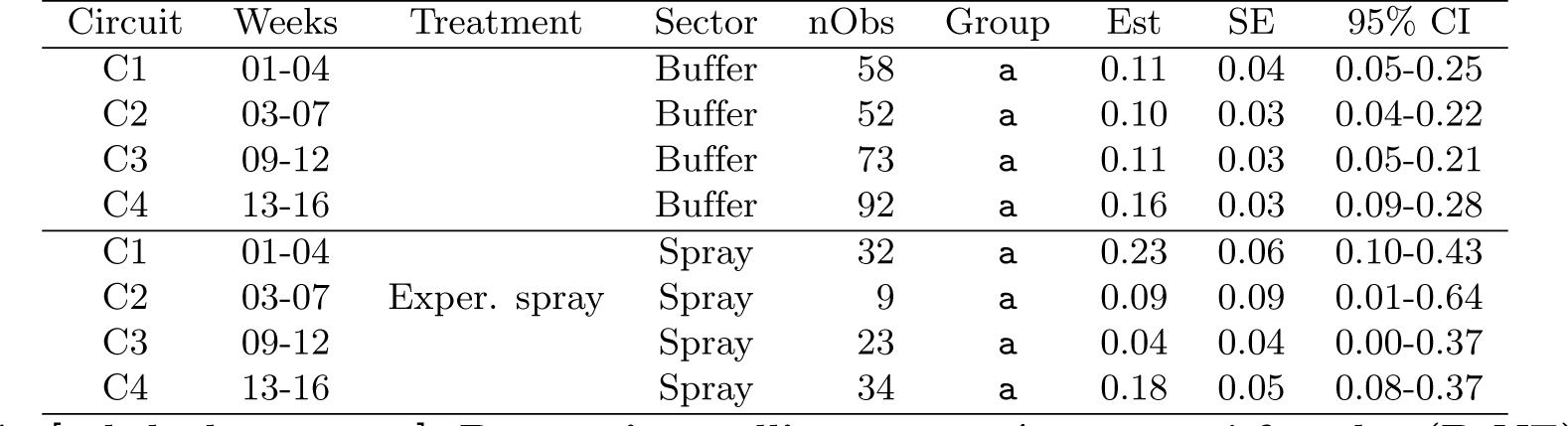
Proportion nulliparouous *Ae. aegypti* females (PrNF), 2013. Model estimates by circuit and treatment sector. Horizontal line separates treatment sectors; significance groups (Tukey HSD) compare among all rows. See also Fig. S5

**Table S8B. [tab hsd par 2014].**
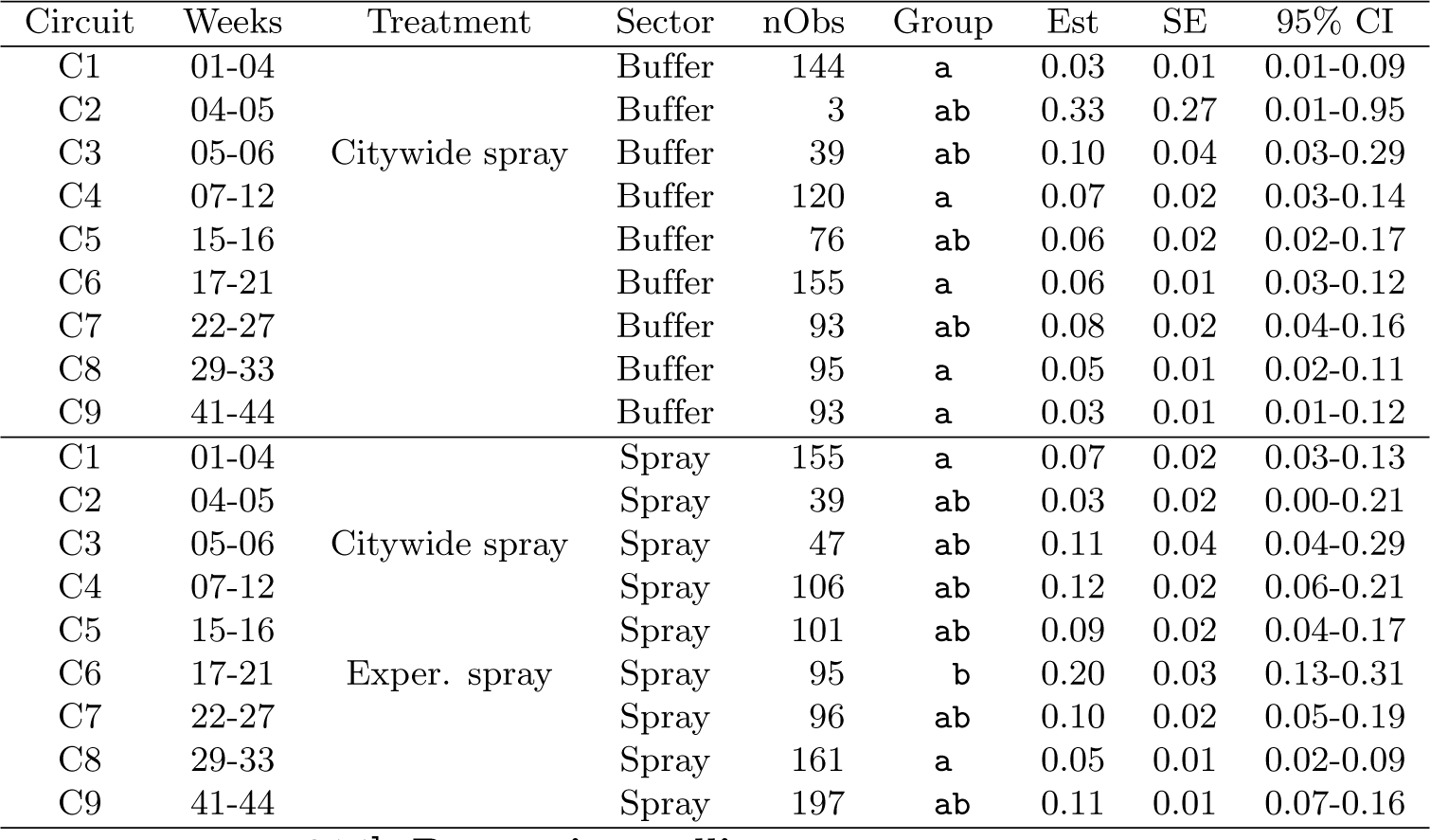
Proportion nulliparouous Ae. aegypti females (PrNF), 2014. See Table S8A for details

**Table S9A. [tab_hsd_bi_2013].** *Ae. aegypti* Positive Containers per House (PC/HSE), 2013. Note that Breteau index (BI) = 100× estimate. Model estimates by circuit and treatment sector. Horizontal line separates treatment sectors; significance groups (Tukey HSD) compare among all rows. No container surveys were conducted during spraying. See also Fig. S5.

| Circuit | Weeks | Sector | nObs | Group | Est | SE | 95% CI |
| --- | --- | --- | --- | --- | --- | --- | --- |
| C1 | 01-04 | Buffer | 565 | a | 0.09 | 0.02 | 0.060-0.147 |
| C3 | 09-12 | Buffer | 590 | ab | 0.16 | 0.02 | 0.111-0.234 |
| C4 | 13-16 | Buffer | 583 | ab | 0.15 | 0.02 | 0.103-0.221 |
| C1 | 01-04 | Spray | 297 | ab | 0.09 | 0.02 | 0.051-0.175 |
| C3 | 09-12 | Spray | 282 | a | 0.07 | 0.02 | 0.038-0.148 |
| C4 | 13-16 | Spray | 268 | b | 0.22 | 0.04 | 0.134-0.373 |

**Table S9B. [tab_hsd_bi_2014].** *Ae. aegypti* Positive Containers per House (PC/HSE), 2014. Note that Breteau index (BI) = 100x estimate. See Table S9A for details.

| Circuit | Weeks | Sector | nObs | Group | Est | SE | 95% CI |
| --- | --- | --- | --- | --- | --- | --- | --- |
| C1 | 01-04 | Buffer | 638 | abc | 0.10 | 0.02 | 0.063-0.159 |
| C4 | 07-12 | Buffer | 606 | abc | 0.08 | 0.01 | 0.048-0.131 |
| C7 | 22-27 | Buffer | 514 | a | 0.15 | 0.02 | 0.095-0.237 |
| C8 | 29-33 | Buffer | 629 | bc | 0.06 | 0.01 | 0.034-0.102 |
| C9 | 41-44 | Buffer | 564 | b | 0.04 | 0.01 | 0.023-0.084 |
| C1 | 01-04 | Spray | 649 | abc | 0.10 | 0.02 | 0.064-0.158 |
| C4 | 07-12 | Spray | 710 | abc | 0.08 | 0.01 | 0.052-0.132 |
| C7 | 22-27 | Spray | 613 | a c | 0.12 | 0.02 | 0.076-0.186 |
| C8 | 29-33 | Spray | 621 | bc | 0.06 | 0.01 | 0.037-0.108 |
| C9 | 41-44 | Spray | 551 | abc | 0.08 | 0.01 | 0.045-0.130 |

**Table S10A. [tab_hsd_cont_2013].** Proportion *Ae. aegypti* Positive Containers (PrPC), 2013. Model estimates by circuit and treatment sector. Horizontal line separates treatment sectors; significance groups (Tukey HSD) compare among all rows. No container surveys were conducted during spraying. See also Fig. S5.

| Circuit | Weeks | Sector | nObs | Group | Est | SE | 95% CI |
| --- | --- | --- | --- | --- | --- | --- | --- |
| C1 | 01-04 | Buffer | 565 | a | 0.04 | 0.00 | 0.026-0.053 |
| C3 | 09-12 | Buffer | 590 | b | 0.06 | 0.01 | 0.047-0.079 |
| C4 | 13-16 | Buffer | 583 | ab | 0.06 | 0.01 | 0.045-0.077 |
| C1 | 01-04 | Spray | 297 | ab | 0.04 | 0.01 | 0.023-0.062 |
| C3 | 09-12 | Spray | 282 | ab | 0.04 | 0.01 | 0.024-0.073 |
| C4 | 13-16 | Spray | 268 | c | 0.11 | 0.01 | 0.076-0.145 |

**Table S10B. [tab_hsd_cont_2014].** Proportion *Ae. aegypti* Positive Containers (PrPC), 2014. See Table S10A for details.

| Circuit | Weeks | Sector | nObs | Group | Est | SE | 95% CI |
| --- | --- | --- | --- | --- | --- | --- | --- |
| C1 | 01-04 | Buffer | 638 | abc | 0.05 | 0.01 | 0.033-0.066 |
| C4 | 07-12 | Buffer | 606 | ab | 0.04 | 0.01 | 0.026-0.057 |
| C7 | 22-27 | Buffer | 514 | c | 0.07 | 0.01 | 0.053-0.098 |
| C8 | 29-33 | Buffer | 629 | a | 0.03 | 0.00 | 0.019-0.048 |
| C9 | 41-44 | Buffer | 564 | a | 0.02 | 0.00 | 0.013-0.040 |
| C1 | 01-04 | Spray | 649 | abc | 0.04 | 0.01 | 0.031-0.061 |
| C4 | 07-12 | Spray | 710 | ab | 0.04 | 0.00 | 0.026-0.053 |
| C7 | 22-27 | Spray | 613 | bc | 0.06 | 0.01 | 0.044-0.082 |
| C8 | 29-33 | Spray | 621 | ab | 0.03 | 0.01 | 0.022-0.053 |
| C9 | 41-44 | Spray | 551 | abc | 0.04 | 0.01 | 0.027-0.063 |

